# *Medicago truncatula* Iron-chaperone 1 (ICHAP1) is required for symbiotic nitrogen fixation

**DOI:** 10.64898/2026.03.29.714480

**Authors:** Cristina Navarro-Gómez, Juan Andrés Collantes-García, Mario Rodríguez-Simón, Jiangqi Wen, Hiram Castillo-Michel, Juan Imperial, Viviana Escudero, Manuel González-Guerrero

## Abstract

Hundreds of proteins in the cell require iron (Fe) or Fe-containing cofactors to function. However, how Fe^2+^ or Fe^3+^ are specifically allocated to each of these proteins in plant cells remains largely unknown. It has been proposed that Fe metalation could be driven by specific interactions with Fe-shuttling proteins known as Fe-chaperones. Here, we present the first family of plant Fe^2+^-chaperones (ICHAPs) with orthologues in dicots and monocots. The role of these proteins in Fe distribution to Fe-dependent metabolic processes has been illustrated using symbiotic nitrogen fixation in *Medicago truncatula* root nodules. ICHAP1 is a soluble Fe^2+^-binding protein that interacts with plasma membrane Fe^2+^ transporter NRAMP1, but not with symbiosome Fe^2+^-transporters. *ICHAP1* mutants present altered Fe distribution in cells and they cannot fix nitrogen. A second family member, ICHAP2 is required to target Fe^2+^ to symbiosomes, as it accepts Fe^2+^ from ICHAP1 and interacts with symbiosome Fe^2+^-importer VTL8, but not with NRAMP1. These results indicate a path for Fe^2+^ allocation from the plasma membrane to the symbiosome through specific protein-protein interactions and Fe^2+^ exchange from NRAMP1 to ICHAP1, to ICHAP2, and to VTL8.

## Main

Iron (Fe) is an essential plant nutrient that acts as cofactor of hundreds of proteins involved in almost every physiological process, including energy transduction, photosynthesis, epigenetics, or plant defence^1–3^. However, at slightly higher concentrations, Fe becomes toxic as it can non-enzymatically produce free radicals in Fenton-style reactions, or it can displace other elements from the active site of enzymes^4^. Consequently, cells must maintain a fine balance to ensure adequate Fe levels while preventing toxicity. Substantial efforts have been made to understand Fe uptake and translocation to sink organs (such as shoots) and how this is regulated. As a result, the main transporter families^5–7^, the Fe-chelators^8^, the Fe-sensing ubiquitin ligases^9,10^, and the transcription factors regulating the process^11–13^ have been unveiled. However, the mechanisms ensuring specific Fe allocation to each Fe-protein has not been clearly defined in plants.

Fe is not the only redox active transition metal in the cell that cannot be free in the cytosol. Copper (Cu) trafficking faces the same challenges, and these have been addressed with a group of proteins, Cu-chaperones that act as shuttles^14^. These proteins ferry Cu cofactors from donor to acceptor proteins driven by the specificity of the docking interfaces and not only by the relative metal binding affinities. As a result, specific Cu^+^-chaperones deliver this element from the plasma membrane transporters to Cu,Zn-superoxide dismutases^15^, to mitochondria^16^, or to the secretory pathway^17^. Similarly, Zn-chaperones have recently been unveiled^18,19^. In contrast, cytosolic Fe-chaperones have only been identified in mammals as polyC-binding proteins (PBCPs)^20^, previously only known as RNA-binding proteins^21^. Through K-homology (KH)-domains, PCBPs bind Fe^2+^ with high affinity^22^, accepting it from NRAMP/DMT Fe^2+^ transporters at the plasma membrane and transferring it to Fe^2+^ efflux transporters^23^, to Fe-storage proteins^20^, or direct it for the synthesis of iron-sulfur ([Fe-S]) clusters^24^. Considering the biochemical need to ensure precise Fe allocation, and the evolutive conservation of the major iron homeostasis elements^1,25^, it should be expected Fe-chaperones exist in most other organisms.

Targeted Fe delivery is particularly challenging in legume root nodule cells. In addition to the usual organelle, these cells must allocate large amounts of Fe to new "pseudo-organelles”, the symbiosomes, that host rhizobia differentiated into nitrogen-fixing bacteroids. Within them, 38 Fe atoms are used to synthesize the three different [Fe-S] clusters of each nitrogenase, the only enzyme in the biosphere that can convert N_2_ into NH_3_^26,27^. As a result of symbiotic nitrogen fixation (SNF), nodules may contain up to a third of the total Fe of the organism although they only represent around 5% of the total plant biomass^28^. The nodule large Fe requirement is not only explained by the amount of nitrogenase cofactors, but also by the accessory enzymes synthesizing these cofactors^29–31^, the production of heme-carrying leghemoglobin controlling O_2_ levels in nodules^32^, as well as other enzymes involved in signalling, development, and energy transduction^33,34^. Consequently, while ammonia is obtained from the bacteroids, the host plant provides Fe in addition to photosynthates and other mineral nutrients^27^. SNF reliance on Fe delivery is highlighted by the critical role that this nutrient plays in nodule development, as the key transcriptional factor controlling the process, NIN, regulates and is regulated by Fe levels in the plant^35,36^. As a result, growth on Fe-deficient soils or mutations in Fe-transport systems impair SNF and nodule development^37–40^.

In model legume *Medicago truncatula*, once Fe is introduced into bacteroid containing cells by NRAMP1^40^, it is translocated into the symbiosomes by nodule-specific Fe transporters VTL8 and FPN2^39,41^. Bridging these transporters, we must find Fe-metallochaperone(s) (ICHAP) interacting with NRAMP1, and providing Fe^2+^ to VTL8 or FPN2 through protein-protein interactions. Here we present Medtr7g011080 (ICHAP1) as a candidate Fe^2+^-chaperone. ICHAP1 interacts with NRAMP1, and it binds Fe^2+^ with high affinity. Mutation of *ICHAP1* results in a loss of nitrogenase activity, and altered Fe localization in nodules. While ICHAP1 cannot interact with VTL8 or with FPN2, it is able to transfer Fe^2+^ to ICHAP2 (Medtr2g019090) another Fe^2+^-binding KH-protein that, in turn, interacts with VTL8. These results not only provide evidence of the existence of plant ICHAPs, but they also show the pathway for Fe delivery from the plasma membrane to the symbiosomes in a bucket brigade-like fashion in which Fe^2+^ is handed from NRAMP1 to ICHAP1 to ICHAP2 to VTL8 into the symbiosomes.

## Results

### ICHAP1 is a Fe^2+^-binding protein that interacts with NRAMP1

Considering thar most Fe delivery systems are conserved^25^ and that the KH-domain is present in most organisms^42^, we hypothesized that a putative plant Fe^2+^-chaperone would also carry this domain. As a result, the PCBP1 Fe^2+^-binding KH-domain3 was used to identify similar domains in the *Medicago truncatula* proteome. Among them, 21 were expressed in nodules according to transcriptomic data at the Symbimics database (Table S1). Split-ubiquitin yeast-two-hybrid assays were carried out to identify which among these proteins interacted with plasma membrane Fe^2+^ importer NRAMP1. One positive interaction was observed when the transporter was combined with Medtr7g011080 (renamed ICHAP1) (Fig. 1a). No interaction was observed when ICHAP1 was combined with either FPN2 or VTL8, the *M. truncatula* transporters involved in Fe^2+^ translocation into the symbiosomes (Fig. S1). The interaction between ICHAP1 and NRAMP1 was not an artefact of the yeast-two-hybrid assays, as it could also be observed in BiFC assays in *Nicotiana benthamiana* leaves (Fig. 1b). Protein modelling of ICHAP1 revealed four KH-domains, similar to those of PCBPs proteins. However, only one of these domains had the S/C and D/E amino acids putatively coordinating Fe^2+^ at a distance compatible with iron binding (Fig. 1c)^22^. To confirm this hypothesis, the C-terminal region of ICHAP1 was fused to a Twin Strep tag (ICHAP1_TS_), purified and titrated *versus* metal chelator MagFura-2 (Fig. 1d, e). The competition assay saturated at a ratio of approximately 1:1, consistent with only one KH domain binding Fe. Interestingly, the saturation followed a bimodal binding affinity with a first plateau at a ratio of 0.4 ± 0.1 Fe^2+^. The K_d_ was estimated to be 1.08 ± 0.15 and 1.96 ± 0.07 mM for each of the two Fe^2+^-binding phases. ICHAP1 orthologues were found in all dicot and monocot species tested (Fig. S2). As expected, the model structures of the closest orthologues presented putative Fe^2+^-coordinating sites (Fig. 1f)^43^.

**Fig. 1.**
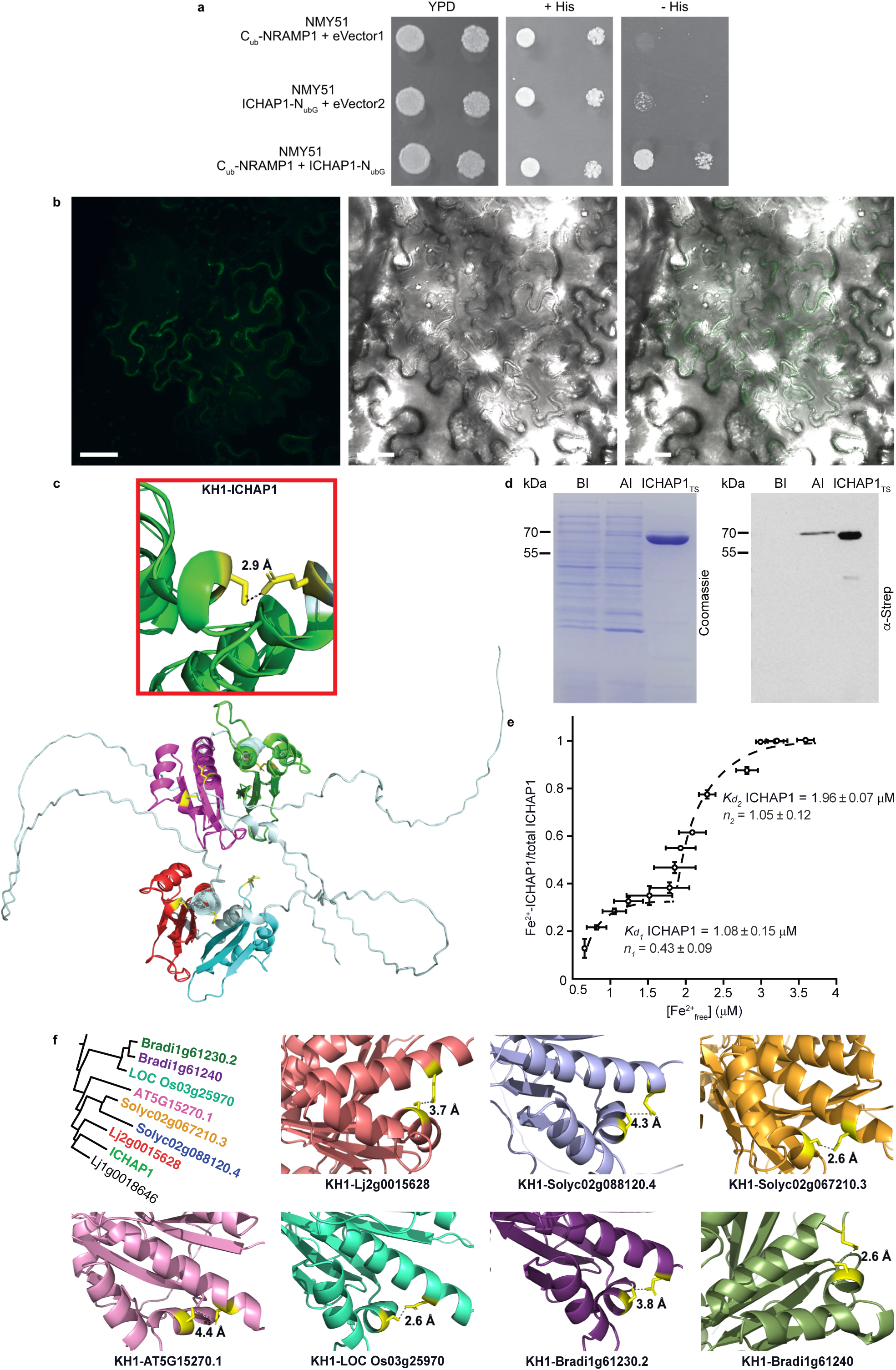
ICHAP1 is a Fe^2+^-binding protein that interacts with Fe^2+^ transporter NRAMP1. **a,** Split-ubiquitin yeast-two-hybrid assay of ICHAP1 and NRAMP1. Top row, yeast containing NRAMP1 fused to the ubiquitin C-fragment at N-terminus (C_ub_-NRAMP1) in pBT3N vector, and empty vector pPR3C (eVector 1); middle row, yeast co-transformed with ICHAP1 fused to modified ubiquitin N-fragment at C-terminus (ICHAP1-N_ubG_) in pPR3C vector, and empty pBT3N (eVector 2); bottom row, yeast co-expressing C_ub_-NRAMP1 and ICHAP1-N_ubG_. Yeasts were grown in YPD and SD +His media as controls, and in SD without histidine (-His) to test the interaction. **b,** BiFC assay of ICHAP1 and NRAMP1. Transient co-expression of ICHAP1 fused to the C-fragment of CFP at C-terminus and NRAMP1 fused to the N-fragment of YFP at N-terminus in *N. benthamiana* leaf cells 3 d-post agroinfiltration. Left panel, fluorescence signal of the positive interaction (green); central panel, transillumination; right panel, overlay of the previous two panels. Bars=50 μm. **c,** Alphafold-modelled ICHAP1 structure. predicted by Alphafold. Its KH domains are shown in green (KH1-ICHAP1), pink (KH2- ICHAP1), cyan (KH3- ICHAP1), and red (KH4- ICHAP1). Boxed inset shows the predicted distance between putative Fe_2+_-coordinating residues S62 and D132 in KH1-ICHAP1. **d,** Purification of ICHAP1_TS_. Left panel, Coomassie Brilliant Blue staining; right panel, immunodetection of *E. coli* BL21 protein extracts before protein induction (BI), after induction (AI), and after Strep purification (ICHAP1_TS_). **e,** Fe^2+^-binding affinity of ICHAP1. Dissociation constants (*K_d1_* and *K_d2_*) of ICHAP1 for Fe^2+^ were determined using Mag-fura-2 competition. Data are means ± SD (n=3). **f,** Predicted KH1 domains of the closest orthologs of ICHAP1 (Fig. S2) from *Lotus japonicus* (Lj2g0015628)*, Solanum lycopersicum* (Solyc02g088120.4 and Solyc02g067210.3), *Arabidopsis thaliana* (AT5G15270.1), *Oryza sativa* (LOC Os03g25970), and *Brachypodium distachyon* (Bradi1g61230.2 and Bradi1g61240). Their candidate Fe^2+^-binding residues are shown in yellow (KH1-Lj2g0015628, S55 and E155; KH1-Solyc02g088120.4, S91 and E143; KH1-Solyc02g067210.3, S56 and D126; KH1-AT5G15270.1, S63 and D133; KH1-LOC Os03g25970, S53 and D123; KH1-Bradi1g61230.2, S53 and D153; KH1-Bradi1g61240, S55 and D125). The predicted distances between these amino acids are indicated.

### ICHAP1 is a cytosolic protein expressed in all plant organs

Studies of *ICHAP1* expression in *M. truncatula* organs indicated that it was expressed in shoots, roots, and nodules (Fig. 2a). No significant changes in expression levels were observed by nodulation. Hairy root transformation with *ICHAP1* promoter region (2 kb upstream of the start codon) controlling the *β*-*glucuronidase* (*gus*) gene confirmed the expression in roots and nodules (Fig. 2b). Cross sections showed that *ICHAP1* was mostly expressed in the nodule core and in the root stele (Fig. 2c, d). Immunolocalization of ICHAP1 fused to three hemagglutinin (HA) tags and expressed under its own promoter confirmed this expression pattern (Fig. 3a, b). At higher magnification, ICHAP1-(HA)_3_ was present among the symbiosomes and in the nuclei, and it was detected in *Sinorhizobium meliloti* colonized and non-colonized cells (Fig. 3c). Transmission electron microscopy using gold-conjugated antibodies confirmed the cytosolic localization of ICHAP1-(HA)_3_ (Fig. 3d), and in some cases, gold particles could also be detected in the proximity of the plasma membrane (Fig. 3e). No specific association of gold particles and symbiosomes was observed (Fig. 3f). To confirm that none of these observations were artefactual controls for autofluorescence of the tagged proteins were carried out (Fig. S3). The specificity of the antibodies used has already been reported for *M. truncatula* nodules^44^.

**Fig. 2.**
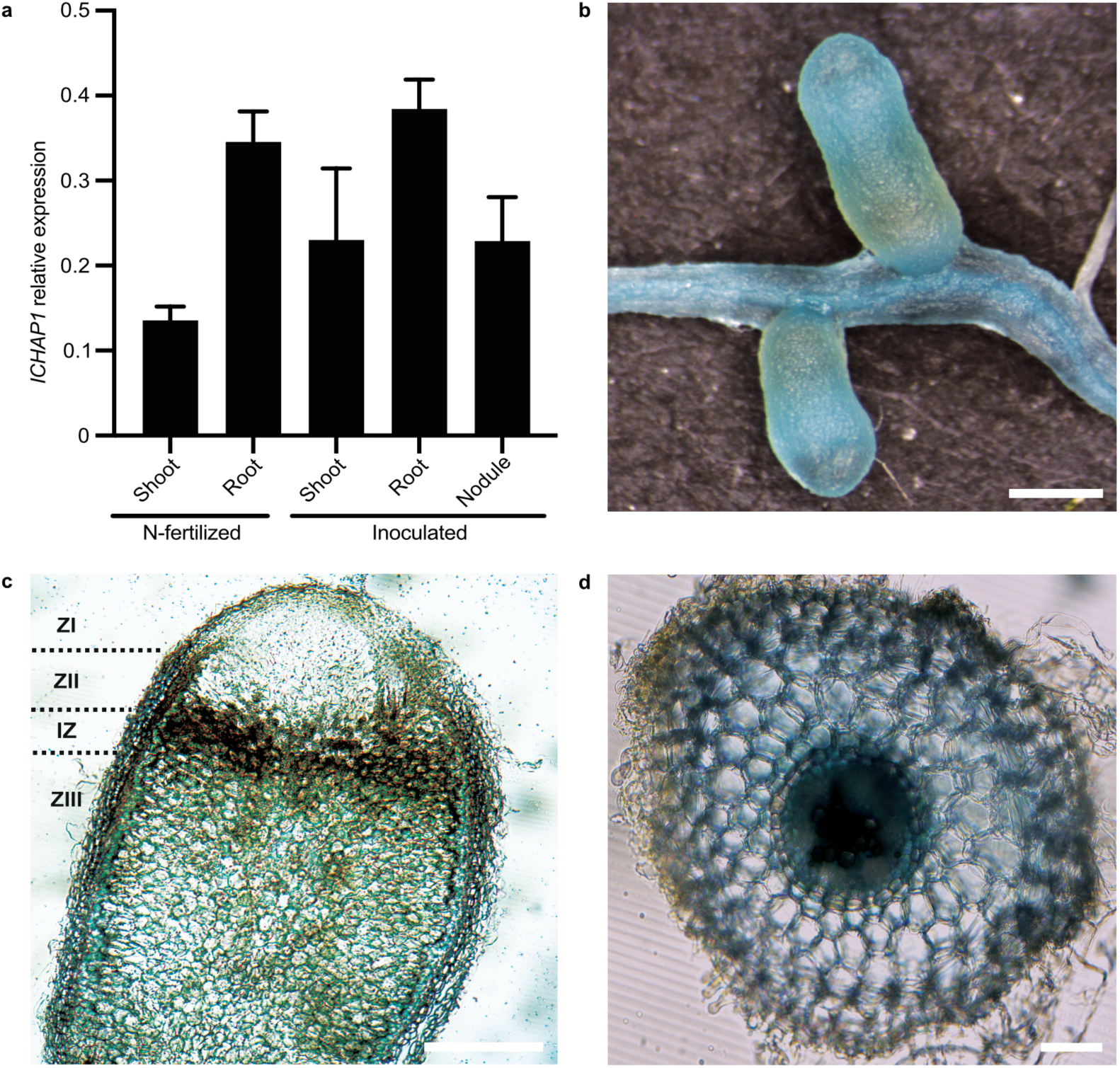
ICHAP1 is expressed in shoots, roots, and nodules. **a,** *ICHAP1* expression in roots and shoots of *M. truncatula* un-inoculated plants and in roots, shoot, and nodules in *S. meliloti*-inoculated ones. Expression is relative to constitutive standard gene *Ubiquitin carboxyl-terminal hydrolase*. Data are the mean ± SEM of three independent experiments with five pooled plants each. **b,** Image of GUS staining of 28-days-post-inoculation (dpi) roots and nodules expressing the *gus* gene under the control of ICHAP1 promoter. Bar=100 μm. **c,** Longitudinal section of a GUS-stained 28-dpi nodule expressing the *gus* gene under the control of ICHAP1 promoter. The different developmental zones (ZI, zone I; ZII, zone II; IZ, interzone; ZIII, zone III) are indicated. Bar=10 μm. **d,** Cross section of a GUS-stained 28-dpi root expressing the *gus* gene under the control of ICHAP1 promoter. Bar=10 μm.

**Fig. 3.**
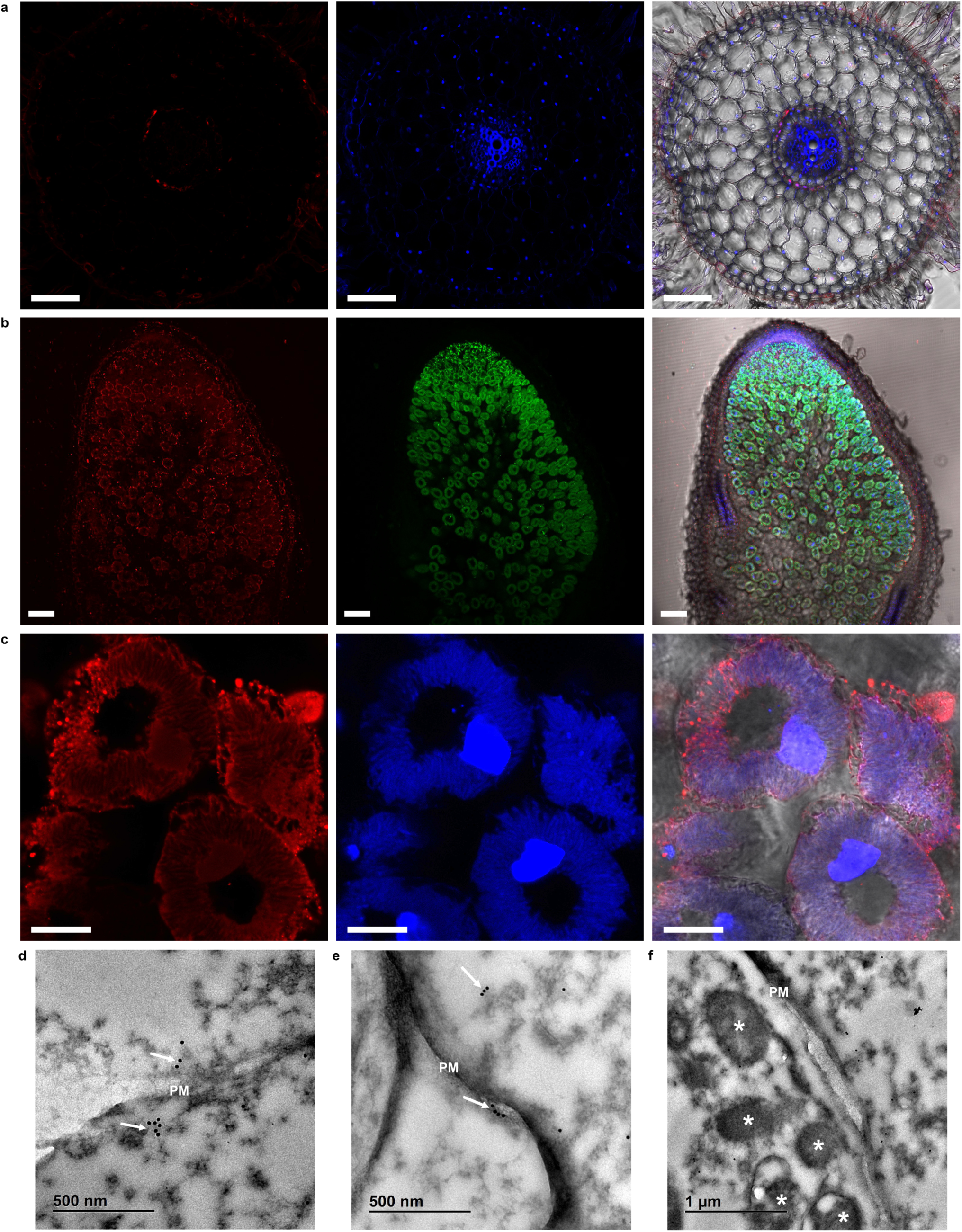
ICHAP1 is a cytosolic protein that can associate to the plasma membrane. **a,** Cross section of a 28-dpi *M. truncatula* root expressing ICHAP1 fused to three HA epitopes (ICHAP-(HA)_3_) driven by its own promoter. Left panel corresponds to the Alexa594-conjugated anti-HA antibody (red), middle panel shows the DAPI-stained root (blue), and right panel is the overlay of the two previous panels with the transillumination. Bars=100 μm. **b,** Longitudinal section of a 28-dpi *M. truncatula* nodule ICHAP-(HA)_3_ driven by its own promoter. Left panel corresponds to the Alexa594-conjugated anti-HA antibody (red), middle panel shows the GFP-labelled *S. meliloti* colonizing the nodule (green), and right panel is the overlay of the two previous panels with the transillumination and the DAPI stain. Bars=100 μm. **c,** Detail of rhizobia-infected cells from the fixation zone of 28-dpi nodule expressing ICHAP-(HA)_3_ under its own promoter. Left panel corresponds to the Alexa594-conjugated anti-HA antibody (red), middle panel shows the DAPI-stained cells (blue), and right panel is the overlay of the two previous panels with the transillumination. Bars=20 μm. **d-f,** Immunolocalization of ICHAP1-(HA)_3_ in 28 dpi nodules using gold-conjugated antibodies and electron transmission microscopy. Clusters of gold particles are indicated by arrows, and asterisks represent bacteroids. PM indicates plasma membrane. Bars=500 nm (d, e), and 1 μm (f).

### ICHAP1 is required for efficient Fe allocation for SNF

Two mutant lines, *ichap1-1* and *ichap1-2,* were obtained with insertions in +269 and +356 leading to a loss of expression (Fig. 4a). When the plants were fertilized with ammonium nitrate and no *S. meliloti* was added, only a mild phenotype was observed. Mutant plants had a small but significant reduction in biomass production, although no significant changes in chlorophyll or iron content were observed (Fig. S4). In contrast, when the plants were inoculated with *S. meliloti* and no fixed nitrogen was provided in the nutrient solution, the mutants presented a reduction on growth and biomass production (Fig. 4b, c). This is likely due to the severe reduction of nitrogenase activity observed in the mutants (Fig. 4d). All these phenotypes were partially restored when a wild-type copy of *ICHAP1* was reintroduced in the mutant background (Fig. S5). None of these phenotypes were complemented when an iron-fortified nutrient solution was used (Fig. S6).

**Fig. 4.**
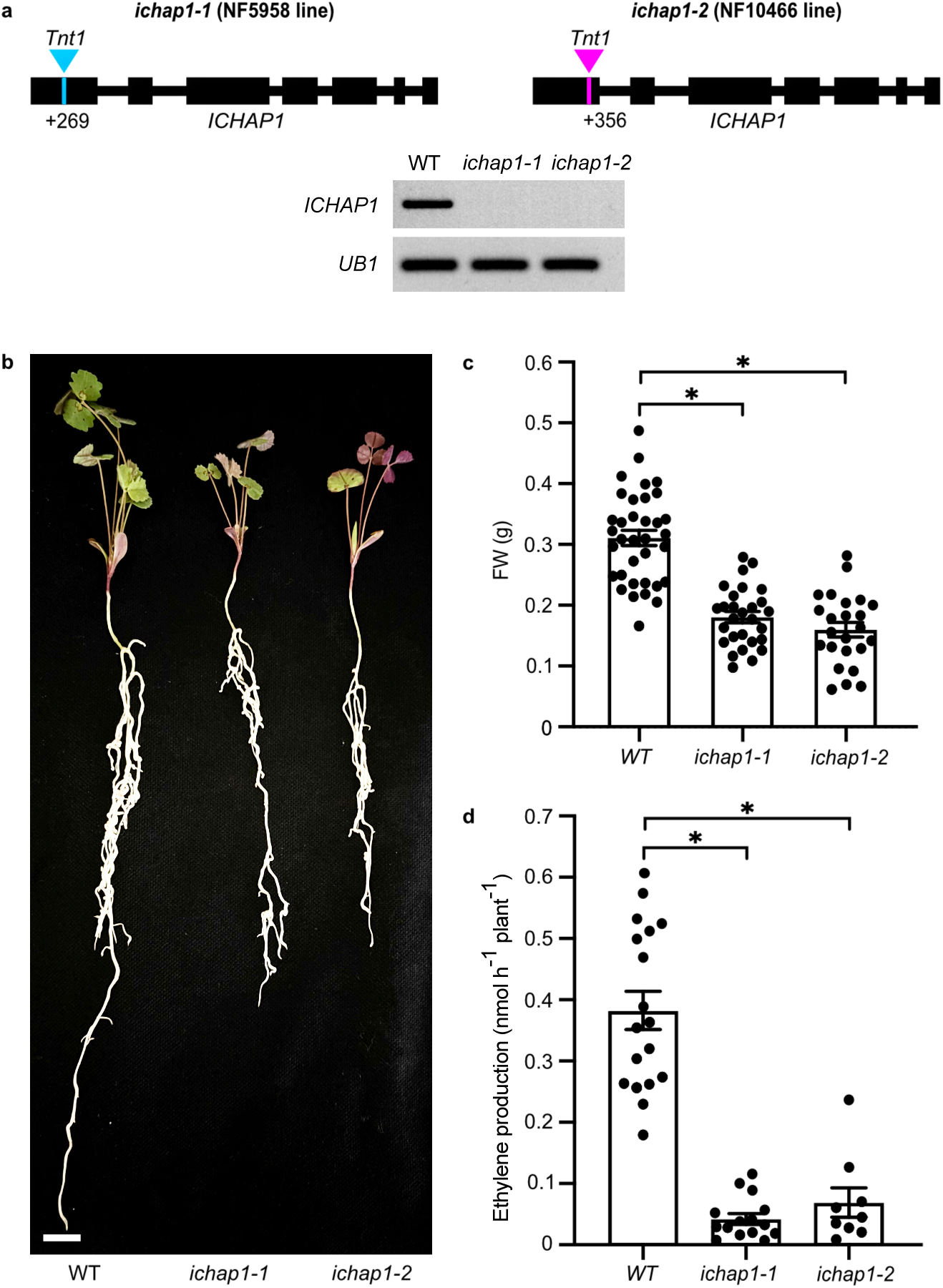
ICHAP1 is required for nitrogen fixation. **a,** Position of *Tnt1* insertion in *ICHAP1* gene for the mutant lines *ichap1-1* (at +269) and *ichap1-2* (at +356). Lower panel, amplification of *ICHAP1* transcript by RT-PCR in 28 dpi roots of wild-type (WT), *ichap1-1* and *ichap1-2 Medicago truncatula* plants. *MtUB1* (*Ubiquitin carboxyl-terminal hydrolase 1*) was used as a constitutive gene. **b,** Growth of representative 28-dpi WT, *ichap1-1* and *ichap1-2* plants. Bar=1 cm. **c,** Fresh weight (FW) of 28-dpi WT, *ichap1-1* and *ichap1-2* plants. Data are the mean ± SEM (n=24-35 pooled plants from three independent experiments). **d,** Acetylene reduction assay in nodules from 28-dpi WT, *ichap1-1* and *ichap1-2* plant*s*. Data are the mean ± SEM (n=10-20 pooled plants from three independent experiments). Two-tailed unpaired Student’s t-test was used for statistical analysis (*, *P* value ≤ 0.05).

Mutant *ichap1-1* nodules had no significant change in coloration or in iron content (Fig S7a, b). Neither were large changes in nodule anatomy observed (Fig. S7c). Iron distribution in wild-type and *ichap1-1* nodules was visualized using synchrotron-based X-ray fluorescence. In the early nodule zones, closest to the apical meristem, no changes were detected (Fig. 5a, b). However, in the late differentiation/early fixation zones, Fe distribution had a completely different pattern for both genotypes (Fig. 5c). Interestingly, while in wild-type nodules iron and sulfur typically colocalized due to the intense synthesis of [Fe-S] clusters, this co-localization was largely lost in *ichap1-1* (Fig. 5d).

**Fig. 5.**
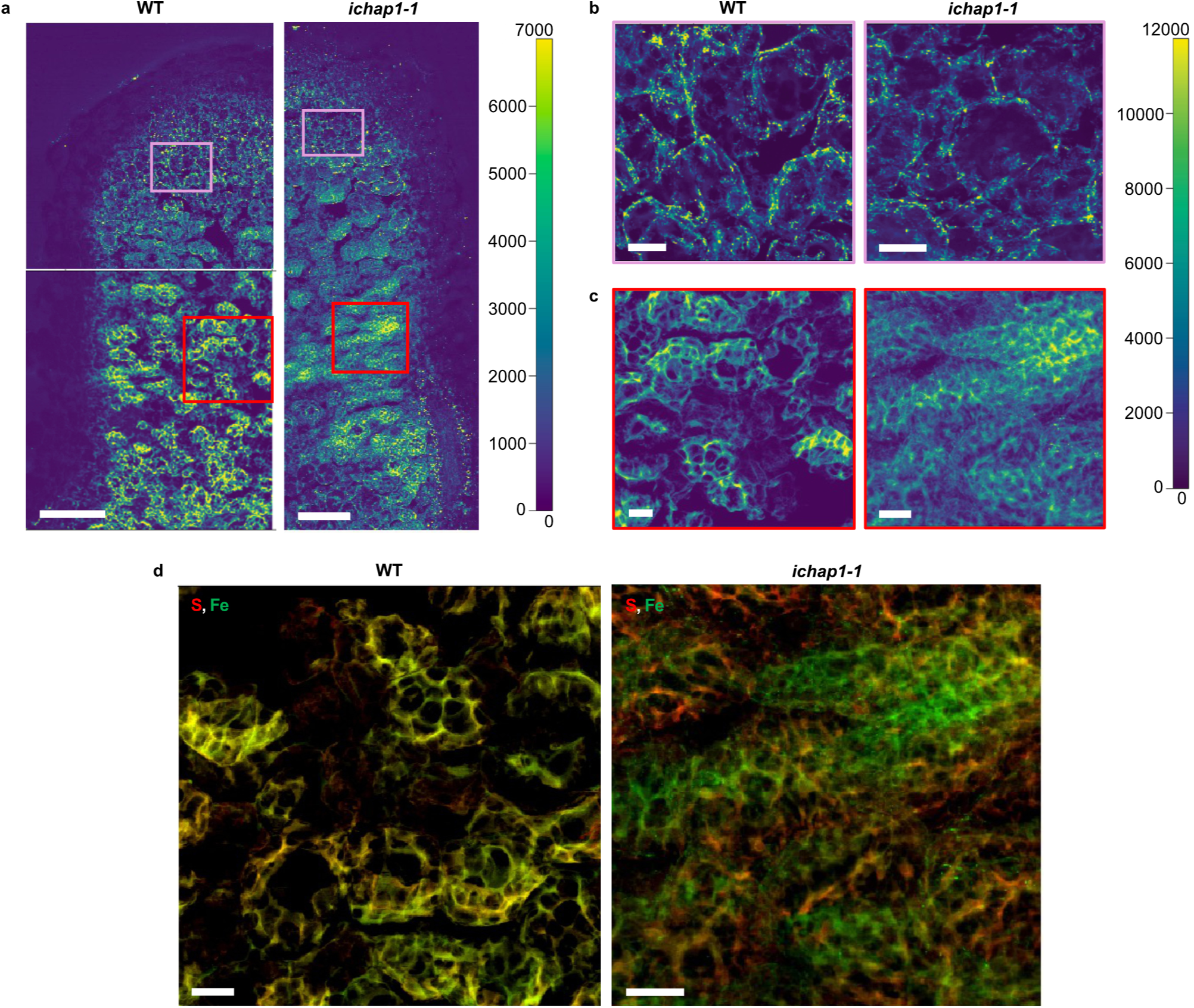
Intracellular Fe distribution is altered in the late differentiation/fixation zone of *ichap1-1* nodules. **a,** Fe distribution in representative wild-type (WT) (left panels) and *ichap1-1* (right panel) nodules. Bars=250 μm. **b,** Higher magnification view of the Fe distribution in the infection/differentiation zone boxed in pink in panel a. Left panel shows WT, and right panel *ichap1-1*. Bars=20 μm. **c,** Higher magnification view of the Fe distribution in the fixation zone boxed in red in panel a. Left panel shows WT, and right panel *ichap1-1*. Bars=20 μm. **d,** Overlay of Fe (green) and S (red) distribution in the fixation zones of WT (left panel) and *ichap1-1* (right panel) nodules. Bars=20 μm. Scale intensity bars indicate normalized fluorescent counts.

### ICHAP1 transfers Fe^2+^ to VTL8-interacting ICHAP2

The interaction of ICHAP1 with NRAMP1, ICHAP Fe^2+^-binding affinity at physiological levels, and the iron-associated phenotype, as well as ICHAP1 cytosolic localization, are indicative of a role on the delivery of Fe from NRAMP1 to Fe-proteins involved in SNF. However, ICHAP1 did not interact with either VTL8 or FPN2 in split-ubiquitin yeast-two-hybrid assays (Fig. S2), nor was it observed associated to symbiosomes (Fig. 3f). Consequently, it is unlikely that this protein is responsible for direct iron allocation to symbiosomes, and at least one other protein should be bridging ICHAP1 and VTL8 or FPN2. To identify this protein, Fe^2+^-bound ICHAP1_TS_ was loaded into a Strep column and nodule extracts were passed through it. Proteomic analyses of the eluted fraction showed Medtr2g019090 (G7IP63), renamed ICHAP2, as the most abundant protein after ICHAP1 (Table 1). Interestingly, ICHAP2 is another KH-domain protein that interacted with VTL8 only when high Fe levels were added to the medium (Fig. S8a). This interaction was validated in BiFC assays in *N. benthamiana* (Fig. S8b). No interaction was observed with NRAMP1 or FPN2 regardless of Fe supplementation in the medium. ICHAP2 was purified and shown to also coordinate one Fe^2+^ (Fig. S8c, d). Titration assays with MagFura-2 revealed the same bi-modal Fe^2+^-binding pattern as ICHAP1 (Fig. S8d). In the first stage, a plateau was reached at a 0.69 ± 0.06 Fe^2+^:ICHAP2 ratio with a K_d1_ of 0.82 ± 0.06 mM, while the second stage was reached at 1.15 ± 0.04 ratio with a K_d2_ of 1.33 ± 0.02 mM. When Fe^2+^-loaded ICHAP1_TS_ was combined with (His)_6_-SUMO-tagged ICHAP2 (_H-SUMO_ICHAP2) and passed through a Strep-column, the two proteins co-eluted (Fig. 6a, Fig. S9a). _H- SUMO_ICHAP2 did not interact on its own with the Strep-resin used (Fig. S9b). To determine whether Fe^2+^ was being transferred, the same assay was repeated but using untagged ICHAP2. After incubating the two proteins and separating them in a Strep column, the iron content of the flow through fraction containing ICHAP2 increased substantially (Fig 6b), although no potentially Fe-loaded ICHAP1_TS_ was observed in that fraction (Fig. 6b, Fig. S9c). When the two proteins were separated by a dialysis membrane to test whether iron could dissociate from ICHAP1_TS_ and reach untagged ICHAP2 by diffusion, no such Fe^2+^ transfer was observed (Fig. 6b).

**Fig. 6.**
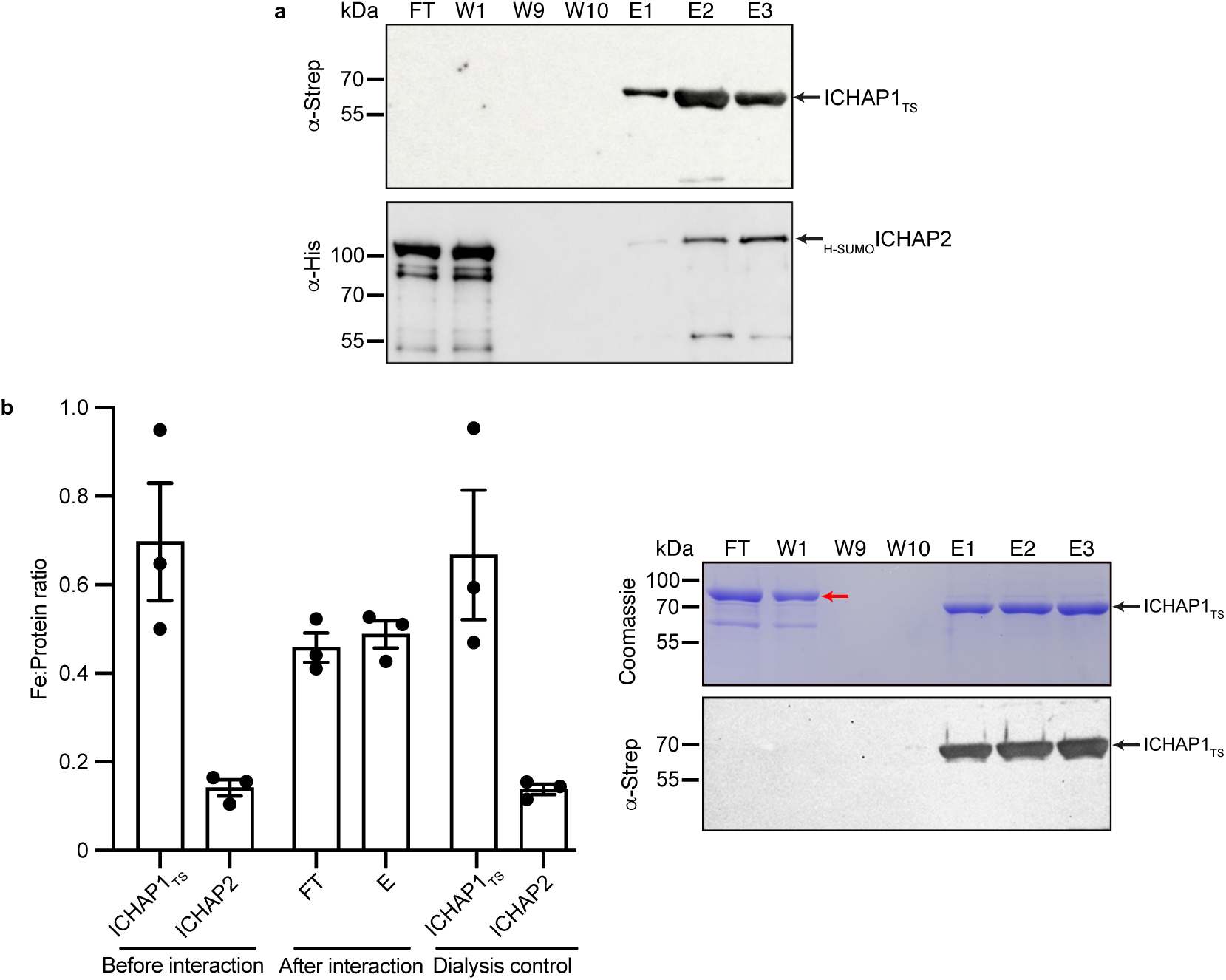
ICHAP1 transfers Fe^2+^ to ICHAP2 by direct protein interaction. **a,** Co-purification of ICHAP1_TS_ and _H-SUMO_ICHAP2 using a Strep-Tactin resin. Upper panel, immunoblot using an anti-Strep antibody; the black arrow indicates ICHAP1_TS_ protein position (62 kDa). Lower panel, immunoblot using an anti-His antibody; the black arrow indicates _H-SUMO_ICHAP2 protein (83 kDa) position. FT corresponds to the flow-through fraction; W1, W9 and W10 to washes 1, 9 and 10; E1-E3, to elutions 1 to 3. Three independent experiments with different pure ICHAP1_TS_ and _H-SUMO_ICHAP2 proteins have been performed. Uncropped membranes are shown in Fig. S9a. **b,** Fe^2+^ transfer from Fe^2+^-loaded ICHAP1_TS_ to ICHAP2. Left panel, molar Fe:protein ratios before the interaction between the two proteins, after the interaction of the two proteins and their separation through a Strep-Tactin column; and dialysis control in which Fe^2+^-loaded ICHAP1_TS_ was separated from ICHAP2 by a dialysis membrane allowing free Fe^2+^ diffusion, but blocking protein transfer from one side to the other. Data are the mean ± SEM of n=3 independent experiments with different pure ICHAP1_TS_ and ICHAP2 proteins. Two-tailed unpaired t-test was used for statistical analysis (*, *P* value ≤ 0.05). Right panel, Coomassie Brilliant blue staining (upper) and anti-Strep immunoblot (lower) of FT, W, and E fractions from the Strep-tag separation of after the interaction between Fe^2+^-loaded ICHAP1_TS_ and ICHAP2 (control for ICHAP_TS_ contamination in the FT fraction). The red arrows indicate the ICHAP2 band. Uncropped gel and membrane are shown in Fig. S9c.

**Table 1.** Top 25 *M. truncatula* proteins co-eluting with ICHAP1.

| Protein ID | Description | Spectral Count |
| --- | --- | --- |
| G7L217 | Putative K domain-containing protein | 2535 |
| G7IP63 | RNA-binding KH domain protein | 1285 |
| G7JLA7 | Alcohol dehydrogenase zinc-binding domain protein | 122 |
| A0A072VE66 | glutamate synthase (NADH) | 119 |
| G7IH71 | phosphoenolpyruvate carboxylase | 117 |
| A0A072TTY6 | Acyl-coenzyme A oxidase | 116 |
| G7KTF6 | Putative tripeptidyl-peptidase II | 113 |
| G7IMW8 | Importin subunit alpha | 83 |
| G7J9B8 | Glutamine synthetase | 75 |
| G7LGQ3 | Putative Acid phosphatase | 74 |
| A0A072VT43 | Translation elongation factor EF-2 subunit | 72 |
| G7L3L0 | 60S ribosomal L8-like protein | 72 |
| A0A072TGQ9 | Alpha-1,4 glucan phosphorylase | 69 |
| G7LIY2 | Lipoxygenase | 63 |
| G7JZK0 | Asparagine synthetase [glutamine-hydrolyzing] | 57 |
| Q84RD9 | Adenosylhomocysteinase | 57 |
| G7JK09 | Polyadenylate-binding protein | 55 |
| Q2HV70 | Proteinase inhibitor I9, subtilisin propeptide | 55 |
| Q1RSH3 | GroEL-like chaperone, ATPase | 54 |
| G7IHV2 | fructokinase | 47 |

## Discussion

As part of heme or [Fe-S] groups, or directly in its ionic forms, Fe is a cofactor of hundreds of different proteins in a plant cell^3^. These proteins are located in the cytosol, in the nuclei, and in the different organelle. From the few plasma membrane transporters, Fe must reach all these proteins without ever being in its soluble hydrated form in the cytosol to prevent oxidative damage and mis-metallation. Plants face the added challenge that Fe is frequently a limiting nutrient in most soil types^45,46^ and, consequently, it must be carefully allocated in a hierarchical manner. Further complexity is reached in legume root nodule cells where the Fe-rich symbiosomes carry out nitrogen fixation^28^. Here, we present the first group of plant Fe-chaperones dedicated to deliver Fe to Fe-dependent metabolic processes, by illustrating their role in SNF.

*Medicago truncatula* ICHAP1 is a Fe^2+^-binding, NRAMP1-interacting protein that is required for SNF. Mutation of *ICHAP1* results in a large loss of nitrogenase activity, which is also observed in *NRAMP1, VTL8, FPN2*, *NAS2,* or *YLS3* mutants^39–41,47,48^, all of them also involved in Fe allocation to nodules. However, ICHAP1 is present in the cytosol and has no known enzymatic activity associated to Fe homeostasis, so its role cannot be transport across membranes or synthesizing known Fe-chelators. Despite this, *ichap1-1* plants have altered Fe distribution in the fixation zones in which no overlap was observed with sulphur, a strong indicator of a deficiency in [Fe-S] cluster synthesis. This is consistent with the loss of nitrogenase activity observed, since it relies on three different [Fe-S] clusters^26^, and with a role of Fe allocation to symbiosomes where the nitrogenase clusters are produced. However, this had to be an indirect effect as ICHAP1-HA was not detected around or within the symbiosomes, nor did it interact with symbiosome Fe transporters FPN2 and VTL8. Pull-down assays revealed that ICHAP1 retained another KH-domain protein, ICHAP2, which was validated in co-immunopurification experiments. As ICHAP1, ICHAP2 coordinated one Fe^2+^ ion per monomer, but complementary to it, it interacted with VTL8, but not with NRAMP1. In additional support for a role in Fe homeostasis, ICHAP2 and VTL8 interaction was only observed in split-ubiquitin yeast-two-hybrid experiments when the culture medium was fortified with Fe. In any case their partnership with Fe-transporters, together with ICHAP1 transferring Fe^2+^ to ICHAP2 through direct protein-protein docking allows us to propose a model for Fe transiting from the cell exterior to the symbiosomes (Fig. 7): Fe^2+^ transported by NRAMP1 is delivered to ICHAP1 to transfer it to ICHAP2 as means to reach VTL8, to which the cation is passed in a bucket-brigade fashion. Additionally, both ICHAP1 and ICHAP2 should still be able to provide Fe to other interacting proteins, an aspect that will be analysed in subsequent studies. Preliminary pull-down experiments already indicate a number of candidate acceptor proteins such as lipoxygenase or glutamate synthase, both known to harbour Fe or [FeS] cofactors^49,50^

**Fig. 7.**
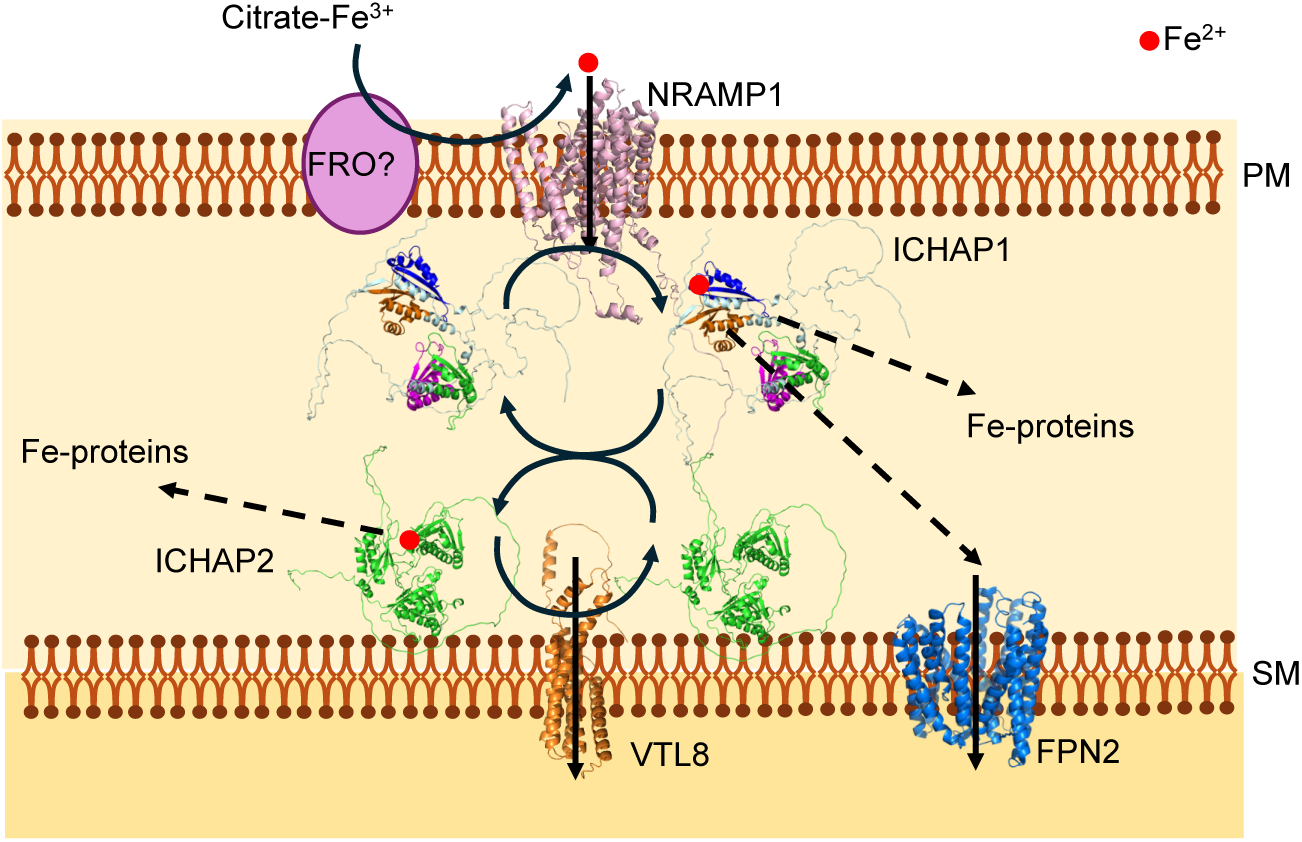
Model of Fe^2+^ transfer to symbiosomes. Apoplastic Fe is bound to citrate as a Fe^3+^-complex. Prior to being transported into the rhizobia-colonized cell by NRAMP1, it must be reduced to Fe^2+^ by a yet-to-be-determined ferroreductase (FRO). ICHAP1 accepts Fe^2+^ from NRAMP1 and transfers it to other Fe-proteins, including ICHAP2, which in turns delivers Fe^2+^ to VTL8 at the symbiosome membranes, and likely to other nodule Fe-proteins. Fe^2+^ delivery to the other symbiosome Fe^2+^-importer, FPN2, is not directly mediated by either ICHAP1 or ICHAP2, suggesting the existence of another nodule ICHAP protein involved in SNF.

However, ICHAP1 is not a nodule-specific protein. It is expressed in all the analysed organs and in all nodule regions, although the strongest phenotype of *ichap1* mutants was detected in nitrogen fixation. Similar observations have been made with mutants in other Fe homeostasis genes that were not nodule-specific, such as *nramp1*, *nas2*, and *ysl3* lines^40,47,48^. This could be explained as the consequence of functional redundancy in roots and shoots, a common phenomenon in plants. In Arabidopsis or rice multiple genes of the same Fe-homeostasis gene family, such as NAS or YSL, need to be simultaneously mutated to produce a strong phenotype ^5,51^, but a single mutant is sufficient to observe strong alterations in legume nodules^47,48^. The most parsimonious explanation to the lack of functional redundancy in nodules is that the evolutionary events that led to the development of these organs occurred at later times than those that caused the expansion of many metal homeostasis gene families. As a result, only one among the members of the Fe homeostasis families acquired competencies in nodules. These observations not only highlight the value of legume root nodules as an excellent model to unveil novel elements in plant metal biology, but they also indicate the existence of additional ICHAP family members in other plant-related processes. This functional redundancy of ICHAP proteins would account for the mild phenotype observed in nitrogen-fertilized *M. truncatula* plants, as well as for the lack of reports on the function of ICHAP orthologues in more usual models, such as *A. thaliana* or rice.

ICHAP1 and ICHAP2 are *bona fide* Fe^2+^-chaperones. Any metallochaperone must interact with either metal transporters or metal donor or accepting proteins, such as Cu^+^-chaperone CopZ with Cu^+^-ATPase CopA^50^, as NRAMP1 with ICHAP1, or as VTL8 with ICHAP2. They must bind their metal with biological relevant affinities, in the nM range for Cu^+^ ^52^, or in the low mM range for Fe as is the case of Fe^2+^-chaperones PCBP1^24^, ICHAP1, or ICHAP2. They must be able to deliver the metal to another apo-metalloprotein through protein-protein interactions^18,20,52^, as it has been shown for Fe^2+^ transfer from ICHAP1 to ICHAP2. Finally, their mutation must affect a metal-dependent process in the cell^45,53^, as nitrogen fixation and Fe distribution are altered by *ICHAP1* mutation. This represents a more controlled metallation pathway than one relying only on a cytosolic labile metal pool following an inverse Irving-Williams series^54^. We could speculate that the role of this pool, integrated by nicotianamine, citrate, glutathione, and other small molecules present in the cytosol^8^, could be serving as a buffer for excess metals, rather than in specific protein metalation.

The metal binding curves of both ICHAPs provide a hint at the iron exchange process. ICHAP1 and 2 present a bimodal Fe-binding model with mid-point saturation at approximately a 0.5:1 Fe^2+^:ICHAP ratio. This is indicative of a situation in which, first, a ICHAP dimer binds Fe^2+^, and once it saturates, it moves to a 1:1 ratio. This possibility of dimerization could reflect the intermediary for Fe^2+^-exchange in which a heterodimer between ICHAP and the acceptor molecule is formed. The existence of a 1:1 ratio ICHAP would reflect a “ready to share” situation in which the availability of Fe^2+^-coordination sites would facilitate docking with the complementary partner. This could be for instance the mechanism of Fe^2+^ delivery from ICHAP1 to ICHAP2. However, even in the “ready-to-share” state, Fe^2+^ bound to ICHAP1 is not easily dissociated, thus preventing the toxic effect of free Fe in the cytosol. As a result, and as it is expected from any metallochaperone, iron transfer only occurs when donor and acceptor proteins are physically interacting with one another. Moreover, the similar binding affinities of ICHAP1 and ICHAP2 could indicate a dynamic scenario of back-and-forth Fe^2+^ exchange in which the direction of Fe fluxes will be mediated by the sink effects of nitrogenase or, conversely, from the recovery of essential limiting Fe once SNF requirements have been fulfilled, or during senescence^55^.

Finally, the discovery of ICHAPs in *M. truncatula* and their orthologues in other plant models opens the way to define a new layer of Fe-homeostasis in plants in which by studying Fe allocation new regulatory processes and new co-evolution partnerships at the docking interfaces between donor and acceptor proteins will be identified. The interactor network of each ICHAPs can be mapped to identify new Fe-proteins, as it is currently being done with Cu^+^-chaperones^44^. Additionally, Fe^2+^-chaperones have the potential to be new tools for biotechnologists for projects of Fe-fortification of crops or to engineer Fe-dependent processes in plants.

## Methods

### Biological materials and growth conditions

*Medicago truncatula* Gaertn R108 seeds were scarified in H_2_SO_4_ for 7.5 min. Then, they were washed with cold water and sterilized with 50 % bleach for 1.5 min and imbibed in sterile water in darkness overnight. Next day, seeds were placed on water-agar plates. After 48 h at 4°C, they were germinated at 22°C for 24 h. Then, seedlings were planted in sterile perlite pots and inoculated with *Sinorhizobium meliloti* 2011 or with *S. meliloti* 2011 expressing pHC60^56^, as indicated. Plants were cultivated in a greenhouse under 16 h light at 25 °C/8 h dark at 22°C conditions, and irrigated with Jenner’s solution or water every 2 d, alternatively^57^. Twenty-eight days post inoculation (dpi), nodules were collected. Non-inoculated plants were grown in similar conditions but watered every 15 d with Jenner’s solution (JS) supplemented with 20 mM NH_4_NO_3_. For metal content experiments, plants were irrigated with a Fe-fortified solution (0.5 g/l Sequestrene 138, Syngenta) every 2 d. For hairy-root transformations, *M. truncatula* seedlings were infected with *Agrobacterium rhizogenes* ARqua1 carrying the appropriate binary vector as described^58^. For agroinfiltration assays, *Nicotiana benthamiana* (tobacco) leaves were infected with *Agrobacterium tumefaciens* GV3101^59^ expressing the interest plasmid constructs. Tobacco plants were grown in a greenhouse, under the same conditions as *M. truncatula*.

*Saccharomyces cerevisiae* strain NMY51 (*MATa, his3Δ200, trp1-901, leu2- 3,112, ade2, LYS2::(lexAop)4-HIS3, ura3::(lexAop)8-LacZ, ade2::(lexAop)8-ADE2, GAL4*) was used for the split-ubiquitin yeast-two-hybrid assays^60^. Yeasts were grown in yeast peptone dextrose (YPD) or in synthetic dextrose (SD) media supplemented with 2% glucose^61^. Protein interaction studies were performed in SD media without histidine.

ICHAP1 and ICHAP2 proteins were produced in *Escherichia coli* strain BL21 (DE3) pLysS (F–*ompT gal dcm lon hsdSB (r_B–_m_B–_)* λ(DE3 [*lacI lacUV5-T7p07 ind1 sam7 nin5*]) [*malB^+^*]_K-12_ (λ^S^) pLysS [T7p20 ori_p15A_](Cm^R^).

### Split-ubiquitin yeast-two-hybrid assay

NRAMP1, VTL8 and FPN2 coding sequences (CDS) were amplified with the primers listed in Table S2 and cloned by homologous recombination in pBT3N vector, that adds to the N-terminal region of each transporter a C-terminal fragment of ubiquitin (C_ub_) fused to a transcriptional activation factor. Similarly, ICHAP1 and ICHAP2 CDSs fragments amplified with the primers listed (Table S2) were cloned by homologous recombination in pPR3C, that adds to the C-terminus the N-terminal fragment of ubiquitin (N_ub_G). A C_ub_-expressing plasmid and a N_ub_G-expressing one were co-transformed in *S. cerevisiae* NMY51. Positive interactors were selected by the recovery of histidine autotrophy caused by the expression of the *HIS3* reporter gene activated by the transcription factor fused to C_ub_.

### Bimolecular Fluorescence Complementation

*NRAMP1* and *VTL8* CDSs were fused at N-terminus to the N-fragment of the Yellow Fluorescence Protein (YFP) in the Gateway vector pNXGW^62^. The CDS of *ICHAP1* and *ICHAP2* were fused to the C-fragment of the Cyan Fluorescence Protein (CFP) at C-terminus in the Gateway vector pXCGW^62^. As negative control, the *gus* gene was cloned also in pXCGW. The primers used for cloning are indicated in Table S2. These constructs were introduced into *A. tumefaciens* GV3101. Agroinfiltration of *N*. *benthamiana* leaves was performed as previously described^63^. Leaves were examined after 3 d by confocal laser-scanning microscopy (Zeiss LSM 880) with excitation light of 488 nm for GFP.

### Protein Expression and Purification

ICHAP1_TS_ was produced by fusing ICHAP1 CDS to two streptavidin (Twin-Strep) tag sequences by using the in-Fusion cloning system (TaKara Bio), in a modified version of pET16b plasmid (courtesy of Dr. Luis Rubio, CSIC, Spain). Primers for the cloning are shown in Table S2. The pET16b ICHAP1_TS_ plasmid was transformed into BL21 (DE3) pLysS *E. coli* competent cells. These cells were grown in 18 l LB media at 37 °C to OD_600_ 0.6, induced with 8 mM lactose and grown for 6 h at 30°C. Cells producing ICHAP1_TS_ (approx. 80 g) were collected and resuspended in 240 ml of buffer W (100 mM Tris pH 7.5, 150 mM NaCl, 10 % glycerol), containing 2.5 mM 1,4-dithiothreitol (DTT), 1 mM Phenylmethanesulfonyl fluoride (PMSF) and 5 μg/ml DNAseI (Sigma). Cells were disrupted by using a French press at 1,500 psi (SLM-AMINCO). The extract was spun down at 54,000 x g for 1 h and loaded onto a 5 ml Strep-Tactin XT 4Flow high capacity column (IBA Lifesciences, Germany). After collecting the flow-through, the column was washed with 20 CV of buffer B and protein was eluted with 7 CV of buffer E (50 mM biotin in Buffer W). The eluted protein was desalted and concentrated using a 30 kDa Amicon centrifugal filter (Amicon Ultra-15, Merck Millipore). Protein purity was analyzed by 12 % SDS-PAGE followed by Coomassie Brilliant Blue staining and Western blot using an anti-Strep MAB antibody (dilution 1:2000; IBA Lifesciences). Protein quantification was performed using the Pierce^TM^ BCA Protein Assay kit (Thermo Scientific).

_SUMO-H_ICHAP2 was the result of cloning ICHAP2 CDS into a pET-His6-SUMO-TEV-LIC cloning vector (2S-T) (Addgene) and transformed into BL21 (DE3) pLysS *E. coli* competent cells. Primers for the cloning are specified in Table S2. Competent cells were grown in 18 l of LB medium at 37 °C until the OD_600_ 0.6 and induced with 1 mM IPTG for 5 h at 30°C. Cells synthesizing _SUMO-H_ICHAP2 (approx. 60 g) were resuspended in 120 ml buffer B (100 mM Tris pH 7.5, 150 mM NaCl, 5 mM imidazole, 10% glycerol) containing 2.5 mM DTT, 1 mM PMSF and 5 μg/ml DNAseI (Sigma), and lysed with a French Press system (SLM-AMINCO). The extract was spun down at 54,000 x g for 1 h and loaded onto a 5 ml IMAC HisTrap HP His-tag protein purification column (Cytiva). Column washing was carried out using a linear gradient starting with buffer B and ending with buffer E (100 mM Tris pH 7.5, 150 mM NaCl, 2.5 mM DTT, 300 mM imidazole, 10% glycerol). _SUMO-H_ICHAP2 protein eluted between 120–300 mM imidazole. Protein was buffer-exchanged with buffer B using 30 kDa Ultra-15 Amicon filtration devices (Merck Milipore). When required, (His)_6_-SUMO tag was cleaved by incubating the protein with His-TEV protease (1:100 TEV:protein ratio) in buffer B with 1 mM tris-2-carboxyethyl-phosphine (TCEP), for 2 h at room temperature. Untagged ICHAP2 was recovered by passing through the nickel column again, in the flow-through and washes. Protein was buffer-exchanged with buffer W using a 30 kDa Ultra-15 Amicon filter (Merck Milipore). Finally, protein purity was analyzed by 12 % SDS-PAGE followed by Coomassie Brilliant Blue staining and Western blot using an anti-His antibody (dilution 1:5000; Sigma).

### Fe^2+^-Binding Affinity determinations

Iron binding affinities of proteins were determined using metal indicator Mag-fura-2 (MF2) (Invitrogen) as described previously^24^. MF2 forms a ratio 1:1 complex with most divalent cations, including Fe^2+^. The apo indicator (MF2_free_) has an absorbance maximum at 366 nm which shifts toward 325 nm concurrent with metal binding. To determine the iron binding affinity of a protein, the decrease in MF2_free_ absorbance at 366 nm was monitored upon iron addition in the presence of the protein. A solution containing 10 μM of either ICHAP1_TS_ or ICHAP2 proteins and 10 μM of MF2 was titrated with 1 mM FeSO_4_ in buffer Z (50 mM HEPES (pH 7.5), 150 mM NaCl, 1 mM TCEP and 10% glycerol). Concentrations of MF2_free_ was determined using extinction coefficients ε_366_ 29,900 M^−1^ cm^−1^ for MF2. Free metal concentrations ([Fe^2+^_free_]) were calculated from *K_d_* = [MF2_free_][Fe^2+^_free_]/ [MF2•Fe^2+^], where K_d_ is the known dissociation constant for MF2 (1.45 μM)^24^. Ratio of metal-bound ICHAP to total ICHAP was plotted as a function of [Fe^2+^_free_] and the data was fitted to a [MF2•Fe^2+^]/[MF2_TOTAL_]= (*n*[Fe^2+^_free_])/(K_d_ + [Fe^2+^_free_]) plot to obtain the protein K_d_ and stoichiometry (*n*). Reported errors for K_d_ and *n* are asymptotic standard errors provided by the fitting software KaleidaGraph (Synergy).

### Quantitative real-time RT-PCR

*Medicago truncatula* RNA isolation and cDNA synthesis were carried out as previously described^40^. Gene expression was analysed by quantitative real-time RT-PCR (9700, Applied Biosystems) with primers indicated in Table S2 and normalized to the *M. truncatula Ubiquitin carboxy-terminal hydrolase* gene (*Medtr4g077320*). Determinations were performed with RNA extracted from three independent biological samples (plant pools), with the threshold cycle determined in triplicate. The relative levels of transcription were determined with the 2^-ΔCt^ method.

### GUS staining

*ICHAP1* promoter region (*ICHAP1p*; 2 kb upstream region of *ICHAP1* start codon) was amplified from *M. truncatula* genomic using the primers indicated in Table S2, and cloned in pDONR207 (Invitrogen) and pGWB3^64^ vectors consecutively by using Gateway Cloning technology (Invitrogen). pGWB3 fused the *β-glucuronidase* (*GUS*) reporter gene to *ICHAP1p*. Hairy-roots transformation was performed as described above. Transformed seedlings were planted in sterile perlite pots and inoculated with *S. meliloti* 2011. GUS activity was measured in nodules of 28 dpi plants as previously described^65^.

### Immunohistochemistry

The *ICHAP1p* and the full-length *ICHAP1* CDS were cloned in tandem (*ICHAP1p::ICHAP1)* in pGWB13^64^ by Gateway technology (Invitrogen). Primers used for this cloning are described in Table S2. This vector fuses three hemagglutinin (HA) epitopes in the C-terminus of the protein (ICHAP1-3xHA). Hairy-root transformation was performed as described above.

For confocal microscopy, transformed seedlings were planted in sterile perlite pots and inoculated with *S. meliloti* 2011 integrating the pHC60 vector, which expresses GFP constitutively. Roots and nodules from 28 dpi plants were collected and fixed in 4 % (w/v) paraformaldehyde, 2.5 % (w/v) sucrose in phosphate-buffered saline (PBS) at 4 °C overnight. Next day, nodules were washed in PBS, and 100 μm sections were generated with a Vibratome 1000 plus (Vibratome). Sections were dehydrated using methanol series (30 %, 50 %, 70 % and 100 % [v/v] in PBS) for 5 min, and then rehydrated. Cell walls were permeabilized with 4 % (w/v) cellulase in PBS for 1 h at room temperature and treated with 0.1 % (v/v) Tween 20 in PBS for 15 min. Sections were blocked with 5 % (w/v) bovine serum albumin (BSA) in PBS and incubated with an anti-HA mouse monoclonal antibody (Sigma) for 2 h at room temperature. After primary antibody, sections were incubated with an Alexa594-conjugated anti-mouse rabbit monoclonal antibody (Sigma) for 1 h at room temperature. After washing the unbound secondary antibody, 4’,6-diamidino-2-phenylindole (DAPI) was used to stain the cell DNA. Images were acquired with a confocal laser-scanning microscope (Leica SP8) using excitation at 488 nm for GFP and at 561 nm for Alexa 594.

The plant material used for electron-microscopy was inoculated with *S. meliloti* 2011. Nodules were collected from 28 dpi plants and fixed in 50 mM potassium phosphate pH 7.4 containing 1 % formaldehyde and 0.5 % glutaraldehyde (fixation solution), for 2 h. Fixation solution was renewed after 1.5 h and left overnight. Samples were washed in 50 mM potassium phosphate pH 7.4 three times during 30 min and three times for 10 min. Then, nodules were dehydrated by incubating with ethanol dilution series of 30 %, 50 %, 70 %, and 90 % during 10 min, 96 % for 30 min, and 100 % during 1 h. Samples were incubated with a series of ethanol and LR-white resin (London Resin Company) dilutions: 1:3 for 3 h, 1:1 overnight, and 3:1 for 3 h. Nodules were included in LR-white resin for 2 d. All incubations were carried out at 4 °C. Nodules were placed in capsules previously filled with LR-white resin and polymerized at 60 °C for 24 h. One-micron thin sections were cut at Centro Nacional de Microscopía Electrónica (Universidad Complutense de Madrid, Spain) with a Reichert Ultracut S-ultramicrotome fitted with a diamond knife. Sections were blocked in 2 % BSA in PBS for 30 min. Then, they were incubated with an anti-HA rabbit monoclonal antibody (Sigma) (dilution 1:20 in PBS) for 2 h. Samples were washed 10 times in PBS for 2 min. As secondary antibody, an anti-rabbit goat antibody conjugated to a 15 nm gold particle (BBI solutions) was used in 1:150 dilution in PBS. Incubation was performed for 1 h. Samples were washed 10 times with PBS and 15 times with water for 2 min, respectively. Sections were stained with 2 % uranyl acetate and visualized in a JEM 1400 electron microscope (JEOL) at 80 kV at Centro Nacional de Microscopía Electrónica.

### Acetylene reduction assay

Nitrogenase activity was analysed by the acetylene reduction assay as described^66^. Nitrogen fixation was determined in mutant and control plants at 28 dpi in 30 ml vials fitted with rubber stoppers. Each vial contained 1-4 plants, depending on the experiment. Three ml of air inside of the vial was replaced by 3 ml of acetylene and incubated at RT for 30 min. Three gas samples of each vial (3 technical replicates of 0.5 ml each) were collected and analysed in a Shimadzu GC-8A gas chromatograph fitted with a Porapak N column (Shimadzu). The amount of ethylene produced was determined by measuring the height of the ethylene peak relative to background.

### Iron content determination

Organ samples (3-20 mg of tissue) were digested with 1 ml of 68 % analytic grade HNO_3_ at 80 °C for 30 min and then allowed to equilibrate at room temperature overnight. Homogenates were diluted with ultra-pure water to a final concentration of 2.5 % HNO_3_ before measurements. In the case of protein samples, they were treated with 68 % analytic grade HNO_3_ at 80 °C for 10 min and then diluted with ultra-pure water to 1 ml (to final 0.68 % HNO_3_). Iron concentrations were determined in an Atomic Absorption Spectrophotometer ContrAA 800 (Analytik Jena) equipped with a graphite furnace, and using commercially available analytic grade metal standards (Inorganic Ventures). All samples were measured in triplicate.

### Synchrotron-based X-ray fluorescence

Micro-X-ray fluorescence (μXRF) hyperspectral images were acquired on the ID21 beamline of the European Synchrotron Radiation Facility (ESRF)^67^, at 110 K in the liquid nitrogen (LN2) cooled cryostat of the new nano scanning X-ray microscope. A nodule from each of at least ten *M. truncatula* R108 and *ichap1-1* plants was embedded into OCT medium and cryofixed by isopentane chilled with LN2. Twenty-five µm sections of frozen samples were obtained using a LN2 cryo-microtome (Leica) and accommodated in an aluminum sample holder cooled with LN2, placed between Ultralene foils (Spex SamplePrep). The beam was focused to 0.18 x 0.2 µm^2^ with a Pt-coated Kirkpatrick–Baez (KB) mirror system providing a photon flux of 2x10^11^ photons s^-1^. The Sample was placed perpendicular to the KB nano-beam. The XRF detection was ensured by two Silicon Drift Diode (SDD) detectors on both sides of the sample, with an angle of 16 degrees with respect to the sample surface. The SDD detectors, one from Mirion (Belgium) and the other one from Rayspec (UK) offered a total active surface of 1.5cm2. A FalconX pulse processor from XIA LLC (US) was used for XRF spectra collection. Images were acquired at a fixed energy of 7.2 keV by raster-scanning the sample in the X-ray focal plane with a step of 2 µm or 0.5 µm and a 50 ms dwell time. Elemental maps were calculated using the PYMCA software package, applying pixel-by-pixel spectral deconvolution to hyperspectral maps normalized by the incoming beam intensity^68^. A semi-transparent beam intensity monitor consisting of a photodiode recording the XRF emission from a Ti coated Si_3_N_4_ window inserted in the beam path, was used for normalization of the measurements by the incoming beam intensity.

### Pull-down assay

30 grams of fresh *M. truncatula* R108 nodules were homogenized with a mortar in buffer A (100 mM Tris pH 7.5, 150 mM NaCl, 100 mM sucrose, 10 % glycerol) with 1 mM PMSF and 5 mg/ml DNAseI (Sigma). The homogenate was centrifuged at 100.000 g 4°C for 1 h (WX-ultracentrifuge, ThermoFisher Scientific). The supernatant containing soluble proteins (“cell free” extract, CFE) was first loaded onto a 0.5 ml Strep-Tactin XT 4Flow High-capacity column (IBA) (control column) to discard unspecific bindings (nodule biotinylated proteins). Then, the resulting flowthrough was loaded into another 0.5 ml Strep-Tactin column previously saturated with Fe^2+^-charged ICHAP_TS_ (ICHAP1 column). ICHAP1 columns were washed with 20 CV of buffer A, and protein elution was carried out with 10 CV of 50 mM biotin in buffer A. Co-eluting proteins with Fe^2+^-ICHAP_TS_ were analysed by 15 % SDS-PAGE, and then identified by Liquid Chromatography Mass Spectrometry (LC-MS/MS) at the Proteomic Service of the Universidad Complutense de Madrid (Spain).

### Protein co-purification assays

15 mM of ICHAP1_TS_ in 2 ml of buffer T (100 mM Tris pH 7.5, 150 mM NaCl, 1 mM TCEP, 10 % glycerol) was bound to 400 μl of Strep-Tactin XT 4Flow high-capacity resin (IBA Lifesciences) by 10 min-batch incubation at room temperature. After packing, the column was washed with 20 CV of buffer T to remove unbound protein. Then, 20 μM of His_6_-SUMO-ICHAP2 in 500 μl of buffer T was loaded onto the Strep-column, and both proteins were incubated together for 10 min at RT to allow the interaction. After collecting the flow-through, the column was washed again with 20 CV of buffer T. The elution was carried out in 1 CV fractions of 50 mM biotin in buffer T. Protein concentrations were determined in all fractions by Pierce^TM^ BCA Protein Assay-Reducing Agent Compatible kit (Thermo Scientific). Protein in selected fractions were visualized by immunoblotting using anti-Strep MAB (dilution 1:2000; IBA Lifesciences) and anti-His (dilution 1:5000; Sigma) antibodies.

### Fe^2+^-transfer assay

ICHAP1_TS_ (15 mM) was incubated for 5 min with 5-fold molar excess of ferrous ammonium sulfate in a final volume of 2.5 ml of buffer T. Metal excess was removed by using a PD10 desalting column (Cytiva). Freshly Fe^2+^-loaded ICHAP1_TS_ was bound to 400 ml of Strep-Tactin XT 4Flow High-capacity resin (IBA Lifescience) by 10 min- batch incubation. After packing, the column was washed with 20 CV of buffer T to remove unbound protein and free Fe. 20 mM of purified ICHAP2 in 500 ml of buffer T containing 5 mM GSH was loaded onto this column, and both proteins were incubated together for 10 min at RT with gentle agitation to initiate Fe^2+^-exchange. The column was washed again with 20 CV of buffer T, and the bound proteins were eluted with 50 mM biotin in buffer T. Protein and iron concentrations were determined in all fractions by Pierce^TM^ BCA Protein Assay-Reducing Agent Compatible kit (Thermo Scientific) and by Atomic Absorption Spectroscopy, respectively. Samples of interest were subjected to Coomassie staining and immunoblot analysis using an anti-Strep MAB antibody (dilution 1:2000; IBA Lifesciences).

### Bioinformatic analyses

To identify KH-domain containing proteins in *M. truncatula* proteome, a BLASTP alignment was performed using the KH3 domain of the *Homo sapiens* PCBP1 (HsPCBP1). As a result, 38 proteins were collected. The analysis was also carried out by using the KH domains of HsPCBP2, but the outcomes were the same, as these two proteins show 83 % of sequence similarity. From this subset, genes which showed expression in both roots and nodules according to Symbimics transcriptomic database (https://iant.toulouse.inra.fr/symbimics)^69^. ICHAP1 orthologues in *Arabidopsis thaliana*, *Solanum lycopersicum*, *Oryza sativa*, *Lotus japonicus*, and *Brachypodium distachyon* were identified by BLASTP analyses (TAIR and Phytozome 14 plant genomic resources) using ICHAP1 full sequence. Protein sequences of the resulting orthologs were obtained from UniProt (http://www.uniprot.org). To build the dendrogram, the Molecular Evolutionary Genetics Analysis (MEGA12) software was used. To identify possible conserved Fe^2+^-binding residues in the orthologues closest to ICHAP1 (Lj2g0015628, Solyc02g088120.4, Solyc02g067210.3, AT5G15270.1, LOC Os03g25970, Bradi1g61230.2, and Bradi1g61240), their annotated KH domains were aligned using Clustal Omega (https://www.ebi.ac.uk/jdispatcher/msa/clustalo) with the predicted ones of ICHAP. The 3D-structures of KH-proteins were modelled by using the Alphafold Server (https://alphafoldserver.com)^70^. Protein visualization and determination of the distance between residues were carried out with PyMOL 3.0.3 (Schörindeger LLC).

### Statistical test

Standard error of the mean (SEM) was calculated, and statistical significance was determined by two-tailed unpaired t-test analysis using Prism7 (GraphPad) software. Test results with *P*-values < 0.05 were considered as statistically significant.

## Acknowledgements

This research was supported by grants PID2021-12460OB-100 and PID2024-155774OB-100 from the Ministerio de Ciencia, Innovación, y Universidades/Agencia Estatal de Investigación/10.13039/50110001103 and “ERDF A way of making Europe”, and UPM grant 200050029 to MG-G. Development of the *M. truncatula Tnt1* mutant population was in part funded by National Science Foundation, USA proposals DBI-0703285, IOS-1127155, and DBI-2233714 to JW. X-ray experiments were carried out in experiment EV-623 on beamline ID21 at the European Synchrotron Radiation Facility (ESRF), Grenoble, France. CN-G and JAC-G were supported by Formación de Personal de Investigación fellowships PRE2019-089164 and PRE2022-101253, respectively, funded by Ministerio de Ciencia, Innovación, y Universidades/Agencia Estatal de Investigación/10.13039/50110001103 and FSE+. We are grateful to Dr. Qingnan Chu (CBGP, UPM-INIA/CSIC) for his technical contributions in assisting in mutant genotyping, and the construction of ICHAP2 expression vector. We would also like to acknowledge other members of laboratory 279 at Centro de Biotecnología y Genómica de Plantas (UPM-INIA/CSIC) for their support and feedback in preparing this manuscript.

## Contributions

CN-G performed most of the experimental work of this manuscript. JAC-G performed ICHAP2 Fe^2+^-binding affinity assays and assisted in the pull-down assays, and together with MR-S provided protein for the biochemical analyses of ICHAP1 and ICHAP2 interactions. JW produced and characterized the *M. truncatula Tnt1* mutant collect. HC-M generated the X-ray fluorescence elemental distribution maps. JI, VE, and MG-G analysed data. VE and MG-G were responsible for overall research supervision and prepared the manuscript with input from all other authors.

## Data availability

All the data used are included in this manuscript.

