## Supporting Figure Legends for "*Medicago truncatula* Iron-chaperone 1 (ICHAP1) is required for symbiotic nitrogen fixation"

### SUPPLEMENTAL FIGURE LEGENDS

**Fig. S1. ICHAP1 does not interact with symbiosome $Fe^{2+}$ transporters VTL8 and FPN2. a**, Split-ubiquitin yeast-two-hybrid assay of ICHAP1-$N_{ubG}$ with $C_{ub}$-NRAMP1. **b,** Split-ubiquitin yeast-two-hybrid assay of ICHAP1-$N_{ubG}$ with $C_{ub}$-VTL8. **c,** Split-ubiquitin yeast-two-hybrid assay of ICHAP1-$N_{ubG}$ with $C_{ub}$-FPN2. Yeasts were grown in YPD and in both standard and 2.5 mM Fe-supplemented SD media with histidine as controls, and in SD media without histidine and supplemented with 2.5 mM $FeSO_4$ to test the interaction. eVector 1: empty pPR3C; eVector 2: empty pBT3N. **d,** Negative control of ICHAP1 and NRAMP1 BiFC assay in *N. benthamiana* leaves 3 d-post agroinfiltration. Co-agroinfiltration of NRAMP1 fused to the N-fragment of YFP at N-terminus and GUS gene fused to the C-fragment of CYP at C-terminus. Left panel shows where fluorescence signal should appear in case of interaction (green); central panel, transillumination; right panel, overlay of fluorescence and transillumination images. Bars=50 μm.

**Fig. S2. Dendrogram of optimally aligned KH-domain proteins orthologous to ICHAP1 in *Arabidopsis thaliana* (AT), *Solanum lycopersicum* (Solyc), *Oryza sativa* (LOC Os), *Lotus japonicus* (Lj), and *Brachypodium distachyon* (Bradi).** ICHAP1 is highlighted by a green arrow.

**Fig. S3. Controls for immunolocalization. a,** Confocal images of representative cross section of un-transformed 28 dpi *M. truncatula* root incubated with both primary anti-HA and secondary Alexa 594-conjugated antibodies. Left panel, HA-tagged proteins detected by using the Alexa 594-conjugated antibody (in red, when positive); central panel, DAPI-stained section (blue); right panel, overlaid images of the transillumination, DAPI-staining, and Alexa 594 channels. Bar=100 μm. **b,** Confocal images of a representative longitudinal section of un-transformed 28 dpi *M. truncatula* nodule incubated with both primary anti-HA and secondary Alexa 594-conjugated antibodies. Left panel, HA-tagged proteins detected by using the Alexa 594-conjugated antibody (in red, when positive); central panel, GFP-expressing *S. meliloti* (green); right panel, overlaid images of the transillumination, DAPI-staining, GFP, and Alexa 594 channels. Bar=100 μm. **c,** Un-transformed 28 dpi *M. truncatula* nodule sections incubated with an anti-HA gold-conjugated antibody. PM, plasma membrane; asterisks, bacteroids. Bars=1 μm.

**Fig. S4. ICHAP1 is not essential for plant growth under non-symbiotic conditions**. **a,** Representative image of non-inoculated wild-type (WT), *ichap1-1*, and *ichap1-2* plants irrigated every two weeks with 2 mM ammonium nitrate. Bar=1 cm. **b,** Fresh weight (FW) of non-inoculated WT, *ichap1-1*, and *ichap1-2* plants irrigated every two weeks with 2 mM ammonium nitrate. Data are the mean ± SEM of n=17-20 plants pooled from three independent experiments. **c,** Chlorophyll content of non-inoculated WT, *ichap1-1*, and *ichap1-2* plants when watered every two weeks with 2 mM ammonium nitrate. Data are the mean ± SEM of n=6 plants pooled from three independent experiments. Two-tailed unpaired t-test was used for statistical analysis (*, *P* value ≤ 0.05). **d,** Iron content of non-inoculated WT, *ichap1-1*, and *ichap1-2* shoots and roots when watered every two weeks with 2 mM ammonium nitrate. Data are the mean ± SEM of n=6-11 plants pooled from three independent experiments. Two-tailed unpaired t-test was used for statistical analysis (*, *P* value ≤ 0.05).

**Fig. S5. ICHAP1 restores the wild-type biomass production and nitrogenase activity in the *ichap1-1* mutant line. a,** Growth of representative wild-type (WT), *ichap1-1* mutant and *ichap1-1* plants transformed with empty vector (WT, *ichap1-1*) or with a vector expressing a wild-type copy of ICHAP1 regulated by its own promoter (*ichap1-1:ICHAP1*). Bar=1 cm. **b,** Fresh weight (FW) of WT, *ichap1-1* and *ichap1-1:ICHAP1* plants. Data are the mean ± SEM (n=10-23 pooled plants from three independent experiments). Two-tailed unpaired t-test was used for statistical analysis (*, *P* value ≤ 0.05). **c,** Acetylene reduction assay in 28-dpi from WT, *ichap1-1* and *ichap1-1:ICHAP1* nodules Data are the mean ± SEM of n=10-23 pooled plants from three independent experiments. Two-tailed unpaired t-test was used for statistical analysis (*, *P* value ≤ 0.05).

**Fig. S6. Iron supplementation does not complement the *ichap1-1* phenotype**. **a,** Growth of representative wild-type (WT) and *ichap1-1* plants watered with standard iron concentrations (Jenner Solution, JS: 7.35 μM Fe) and iron excess (Fe-fortified JS, 0.5 g/l Sequestrene). Bar=1 cm. **b,** Fresh weight (FW) of 28 dpi WT and *ichap1-1* plants watered with JS or Fe-fortified JS. Data are the mean ± SEM (n=9-12 plants pooled from three independent experiments). All comparisons were done to WT samples using two-tailed unpaired student's t-test (*, P ≤ 0.05). **c,** Acetylene reduction assay in 28 dpi WT and *ichap1-1* plants watered with JS or Fe-fortified JS. Data are the mean ± SEM (n=7-12 plants pooled from three independent experiments). All comparisons were done to WT samples using two-tailed unpaired student's t-test (*, P ≤ 0.05).

**Fig. S7. Nodule development and iron content of *ichap1-1* mutant compared to wild-type (WT). a,** Representative images of 28 dpi WT (left panel) and *ichap1-1* (right panel) nodules. Bars=100 μm. **b,** Iron content in 28 dpi shoots, roots and nodules of WT and *ichap1-1* plants watered with the standard Jenner solution (7.35 μM Fe). Data are the mean ± SEM (n=6-9 plants pooled from three independent experiments). Unpaired Student's t-test was used for statistical analysis (*, *P* value ≤ 0.05). **c,** Cross sections of representative 28-dpi WT (left panel) and *ichap1-1* (right panel) nodules. Bars=20 μm.

**Fig. S8. ICHAP2 interacts with symbiosome $Fe^{2+}$-transporter VTL8 in an iron-dependent way. a,** Split-ubiquitin yeast-two-hybrid assays of ICHAP2 and VTL8 (upper panels), NRAMP1 (central panels) and FPN2 (lower panels) $Fe^{2+}$ transporters. For each assay, first row, yeast co-expressing the $Fe^{2+}$ transporter fused to the ubiquitin C-fragment at N-terminus ($C_{ub}$-VTL8/NRAMP1/FPN2) cloned in pBT3N and empty vector pPR3C (eVector 1); second row, yeast co-expressing ICHAP2 fused to modified ubiquitin N-fragment at C-terminus (ICHAP2-$N_{ubG}$) cloned in pPR3C and empty pBT3N (eVector 2); last row, yeast co-expressing $C_{ub}$-VTL8/NRAMP1/FPN2 and ICHAP2-$N_{ubG}$. Yeasts were grown in YPD and in both standard and 2.5 mM $FeSO_4$-supplemented SD media with histidine as controls, and in SD media without histidine and supplemented with 2.5 mM $FeSO_4$ to test the interaction. **b,** BiFC assay of ICHAP2 and VTL8. Upper panels, transient co-expression of ICHAP2 fused to the C-fragment of CFP at C-terminus and VTL8 fused to the N-fragment of YFP at N-terminus in *N. benthamiana* leaf cells 3 d-post agroinfiltration. Lower panels, negative control of ICHAP2 and VTL8 BiFC assay. Co-agroinfiltration of VTL8 fused to the N-fragment of YFP at N-terminus and GUS gene fused to the C-fragment of CYP at C-terminus. Left panels, fluorescence signal of the positive interaction (green); middle panels, transillumination; right panels, overlay of fluorescence and transillumination images. Bars=50 µm. **c,** Purification of $_{\text{H-SUMO}}$ICHAP2 and untagged ICHAP2. Left panel, Coomassie staining; right panel, anti-His immunoblot of *E. coli* BL21 protein extracts before the protein induction (BI), after the induction (AI), the protein after the His purification (Protein Before cleavage, $P_{Bc}$), and after the $His_6$-SUMO tag removal (Protein After cleavage, $P_{Ac}$). The black arrow points the $_{\text{H-SUMO}}$ICHAP2 (83 kDa) and the red, ICHAP2 (69 kDa). **d,** $Fe^{2+}$-binding affinity of ICHAP2. Dissociation constants ($K_{d1}$ and $K_{d2}$) of ICHAP2 for $Fe^{2+}$ were determined using Mag-fura-2 competition. Data are means ± SD (n = 3) independent experiments with different ICHAP2 purifications.

**Fig. S9. ICHAP2 does not bind to the Strep-column. a,** Uncropped immunoblots shown in Fig. 6a. **b,** Immunodetection with an anti-Strep (left panel) and anti-His (right panel) antibodies of $_{\text{H-SUMO}}$ICHAP2 in flowthrough (FT), washes (W1, W9 and W10), and elution (E1-E3) fractions after passing through a Strep-column. **c,** Uncropped immunoblot (left panel) and Coomassie Brilliant Blue staining (right panel) shown in Fig. 6b.
