## Supplementary figures and images for "*Medicago truncatula* Iron-chaperone 1 (ICHAP1) is required for symbiotic nitrogen fixation"

### Supporting Figures

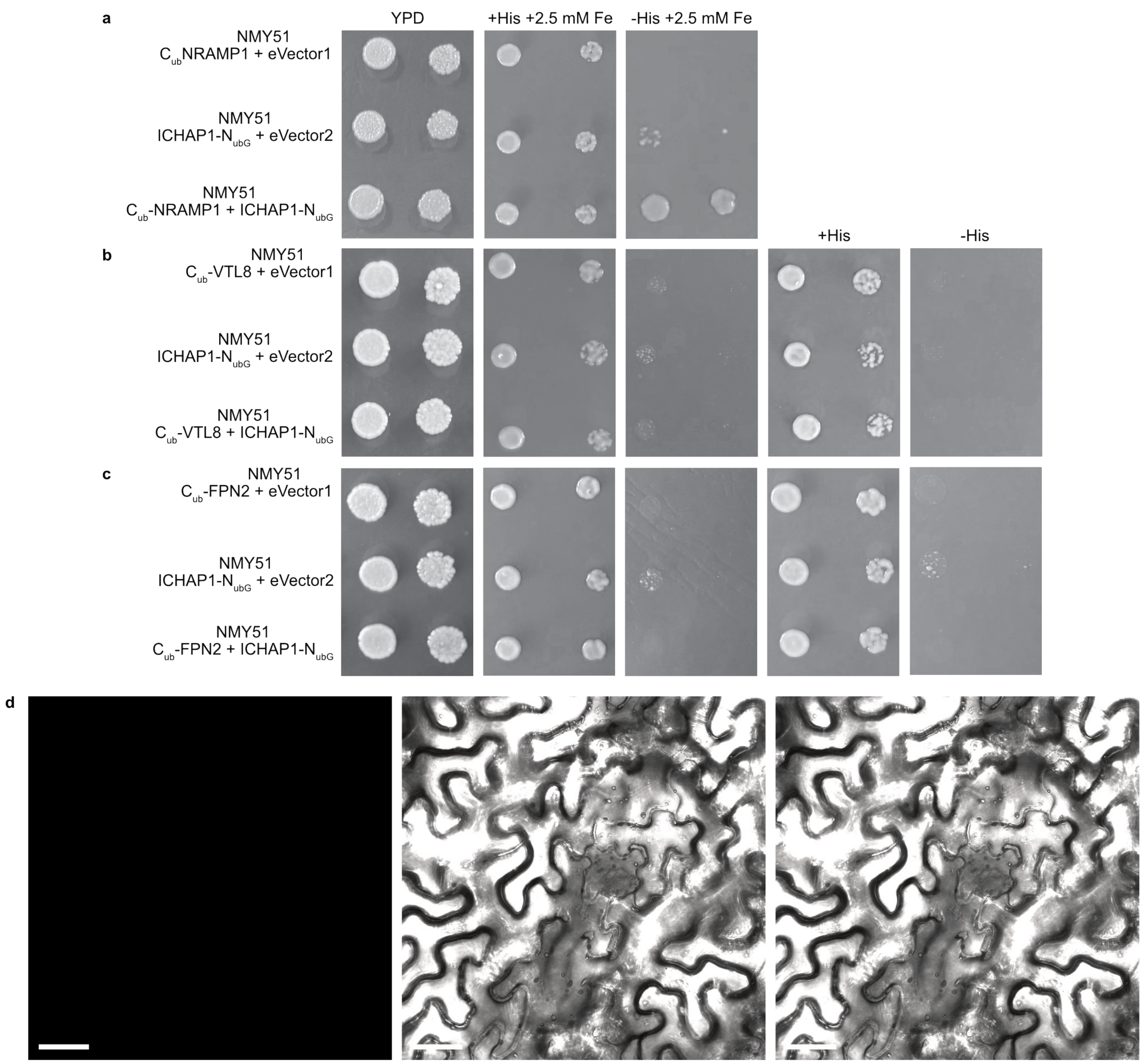

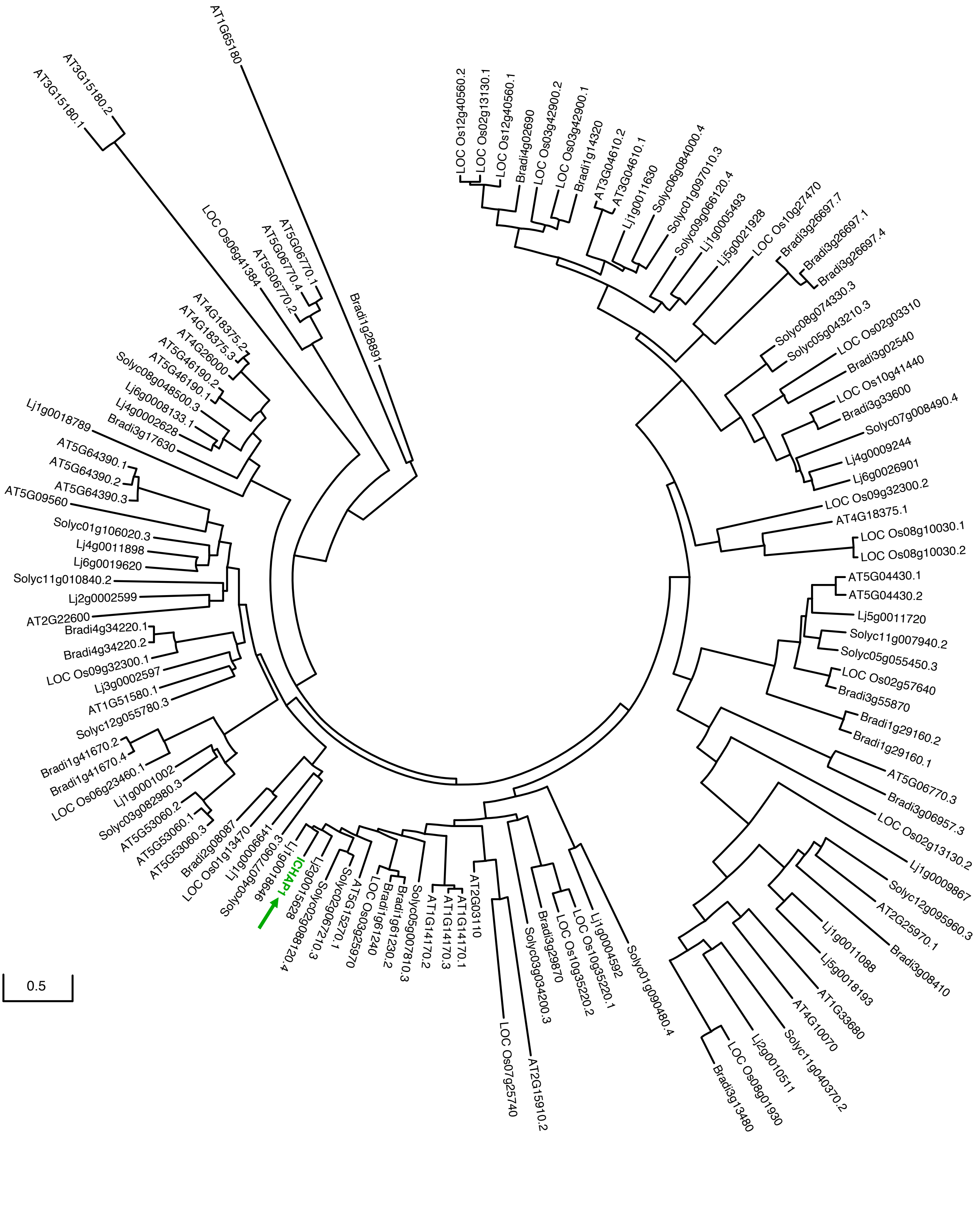

Figure S3

a

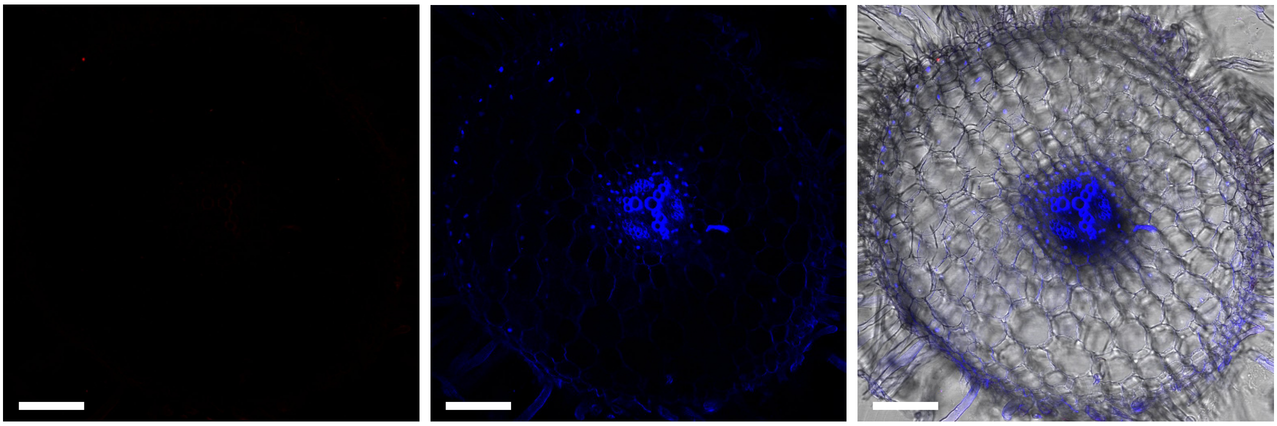

b

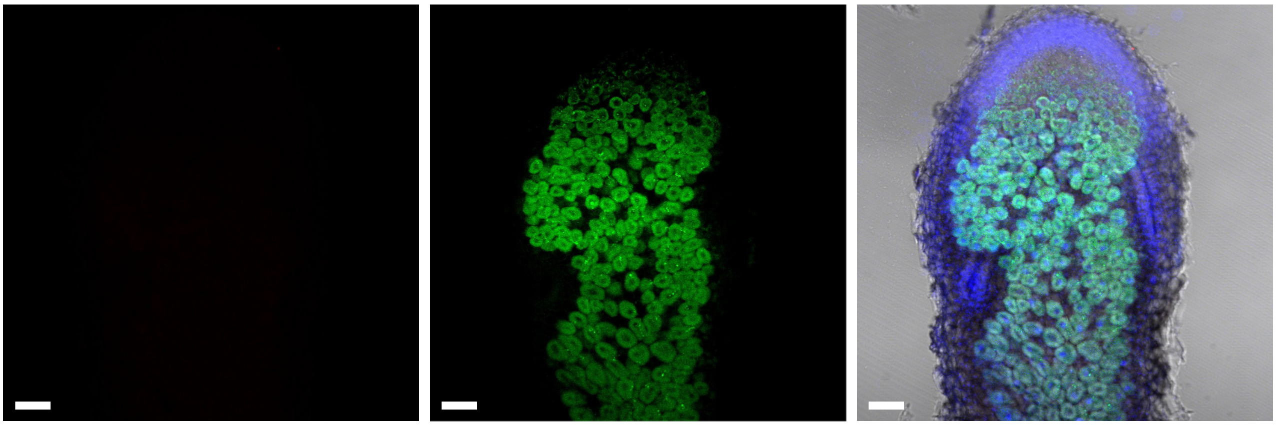

c

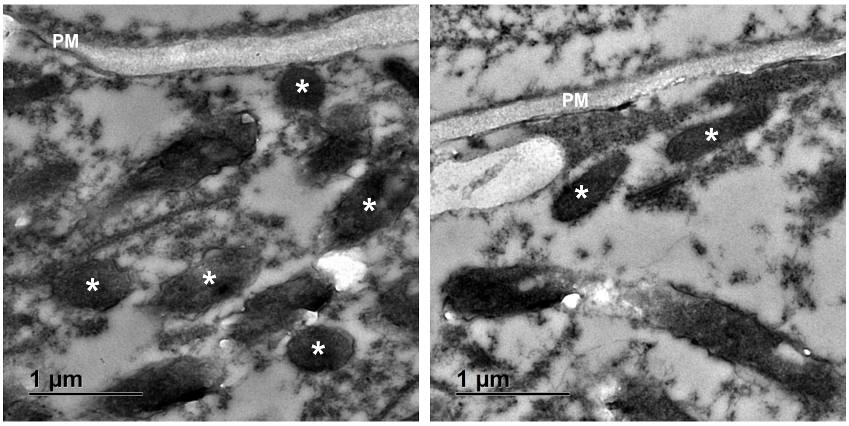

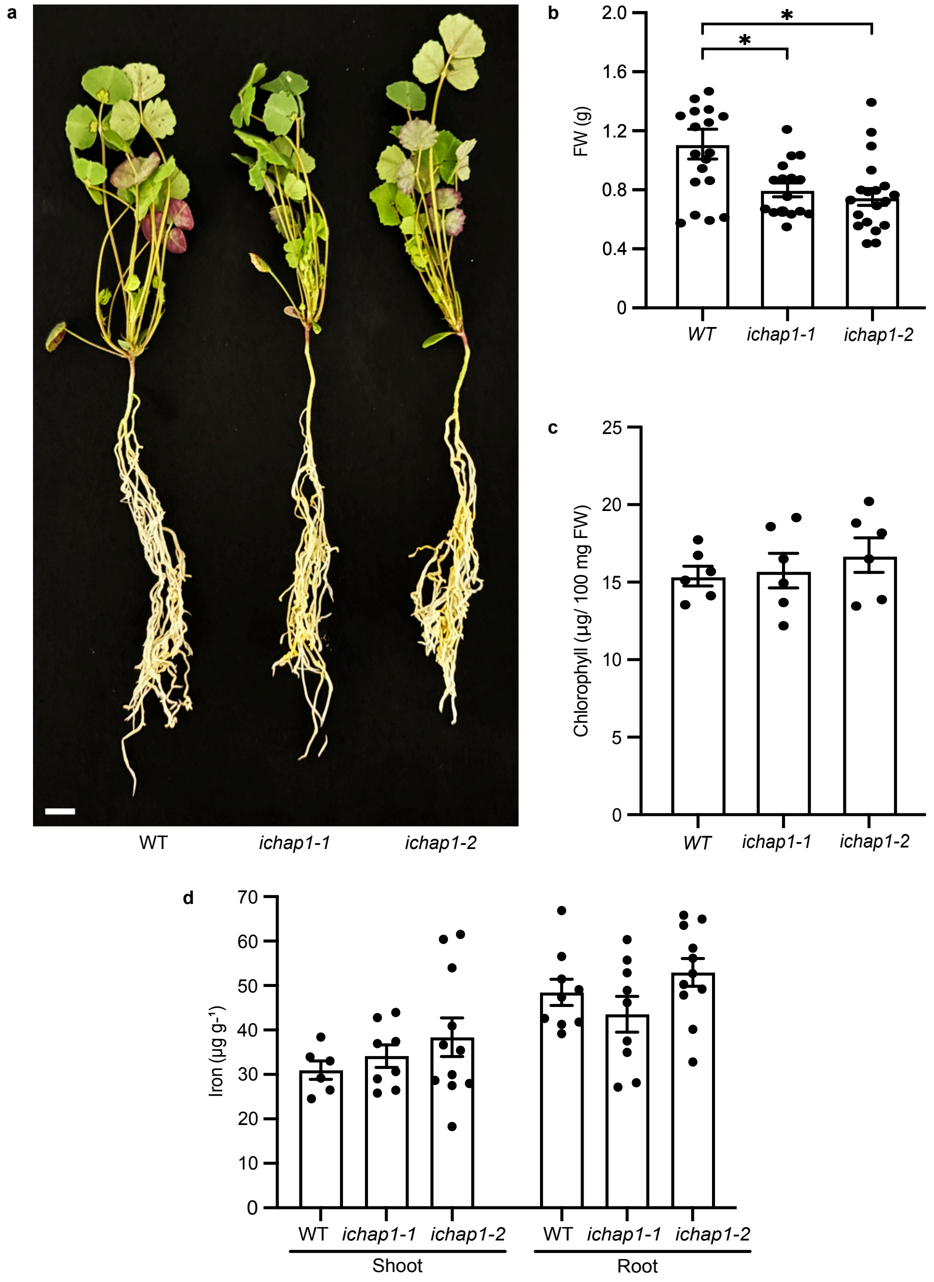

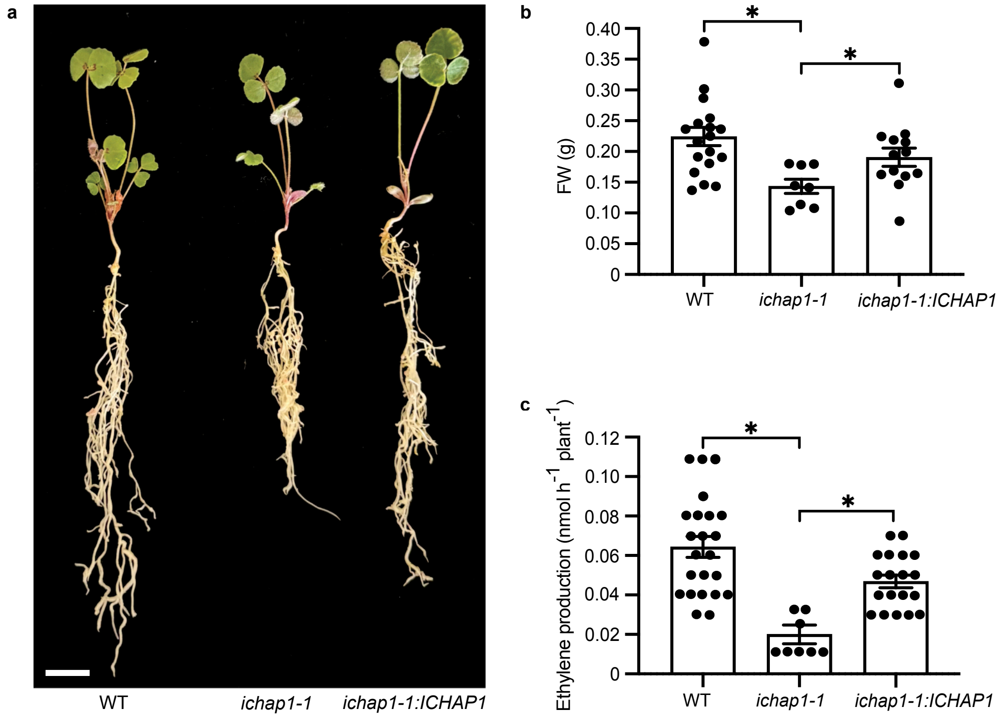

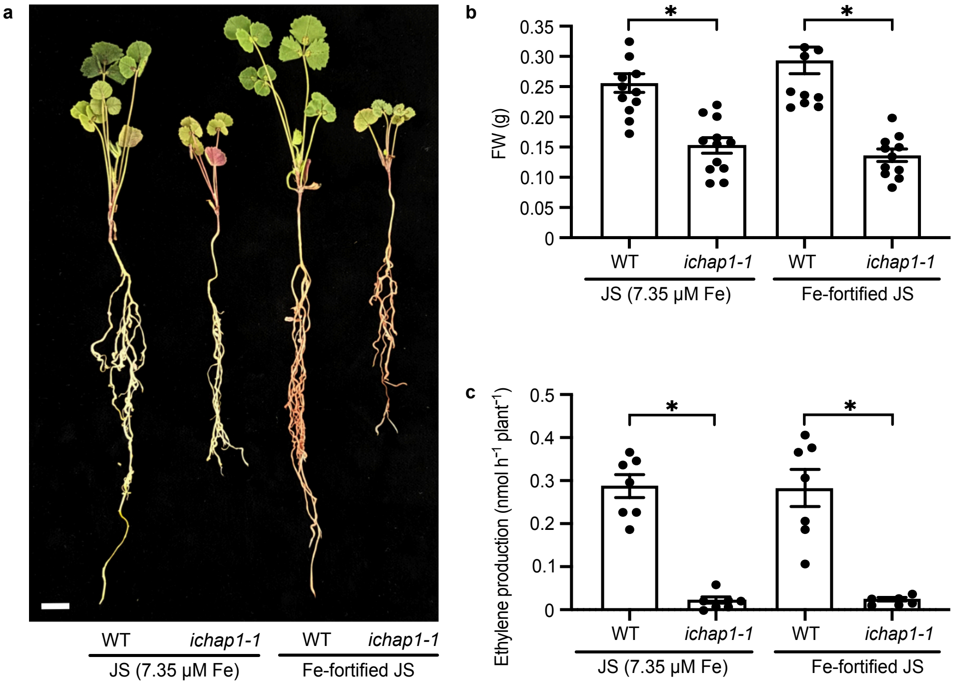

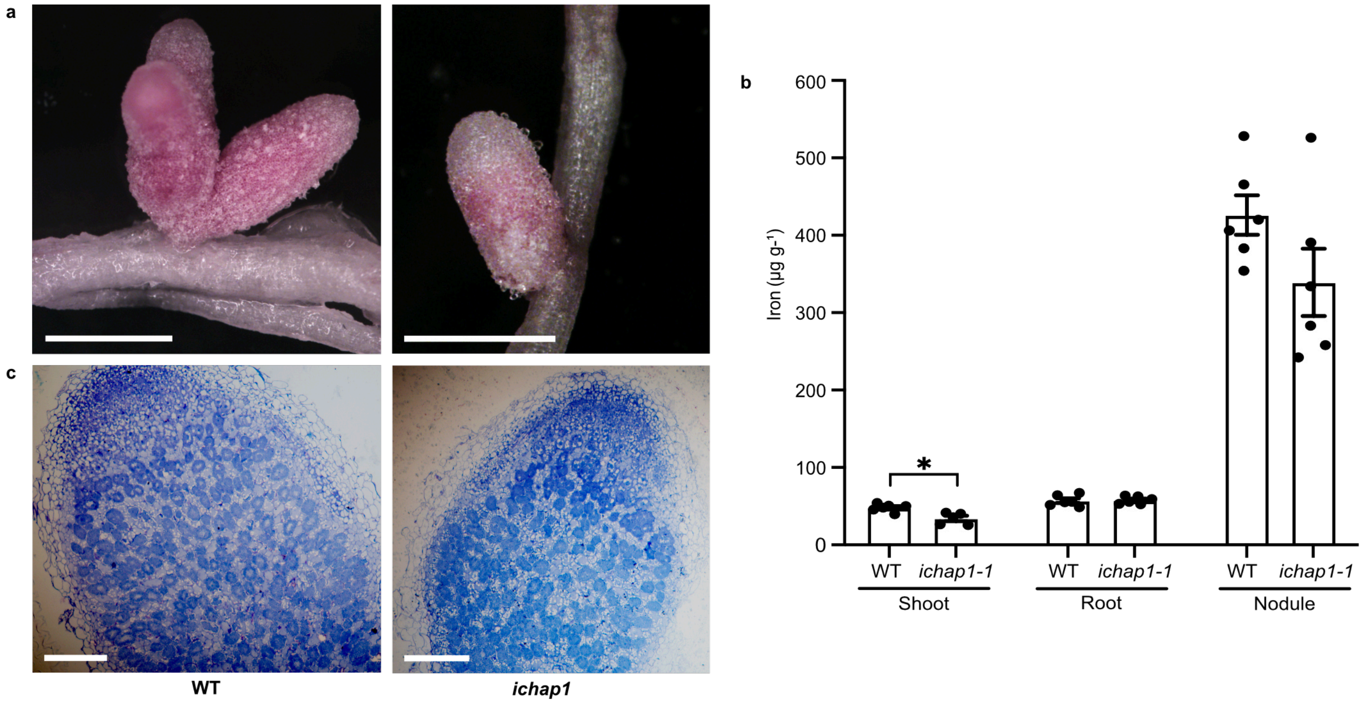

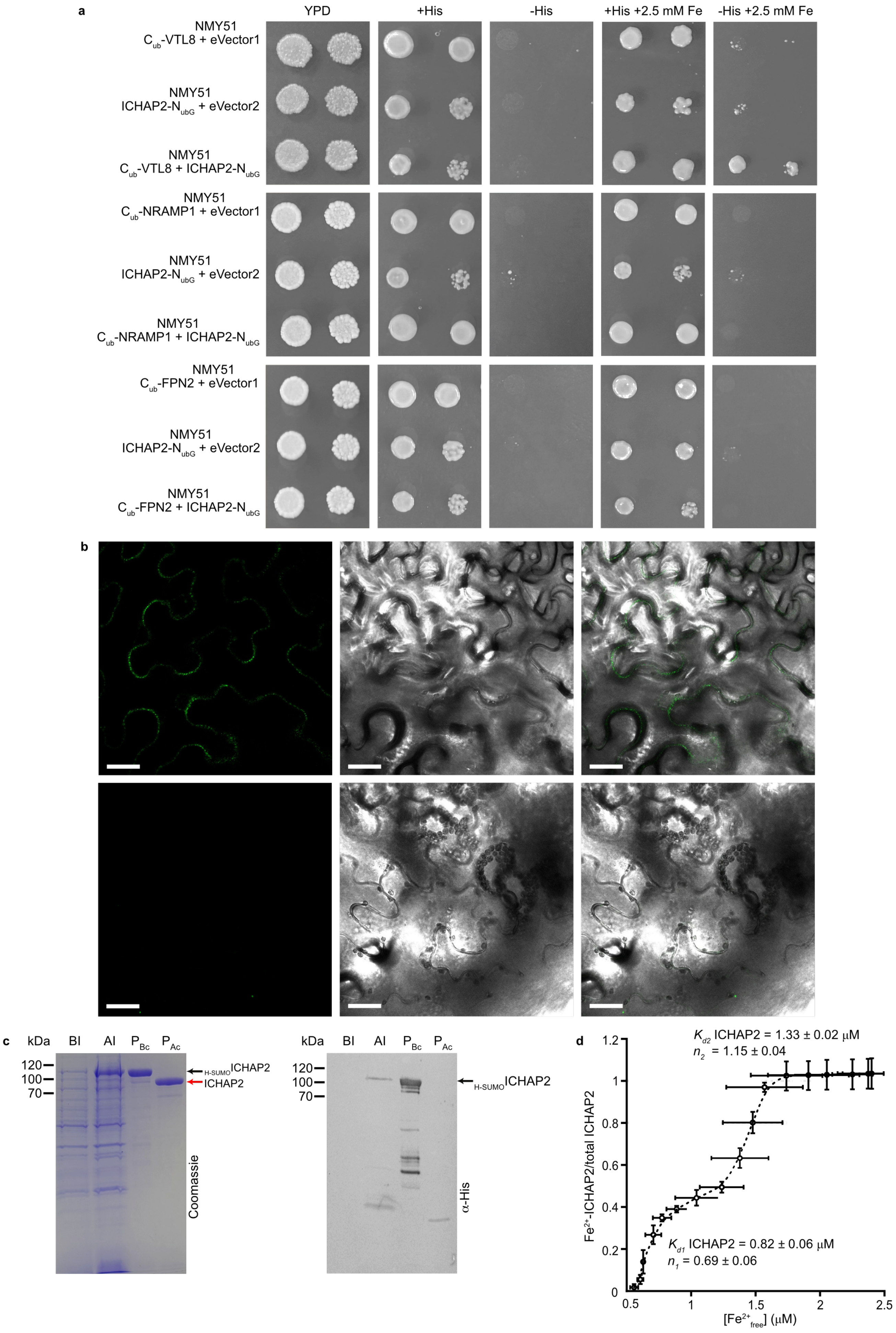

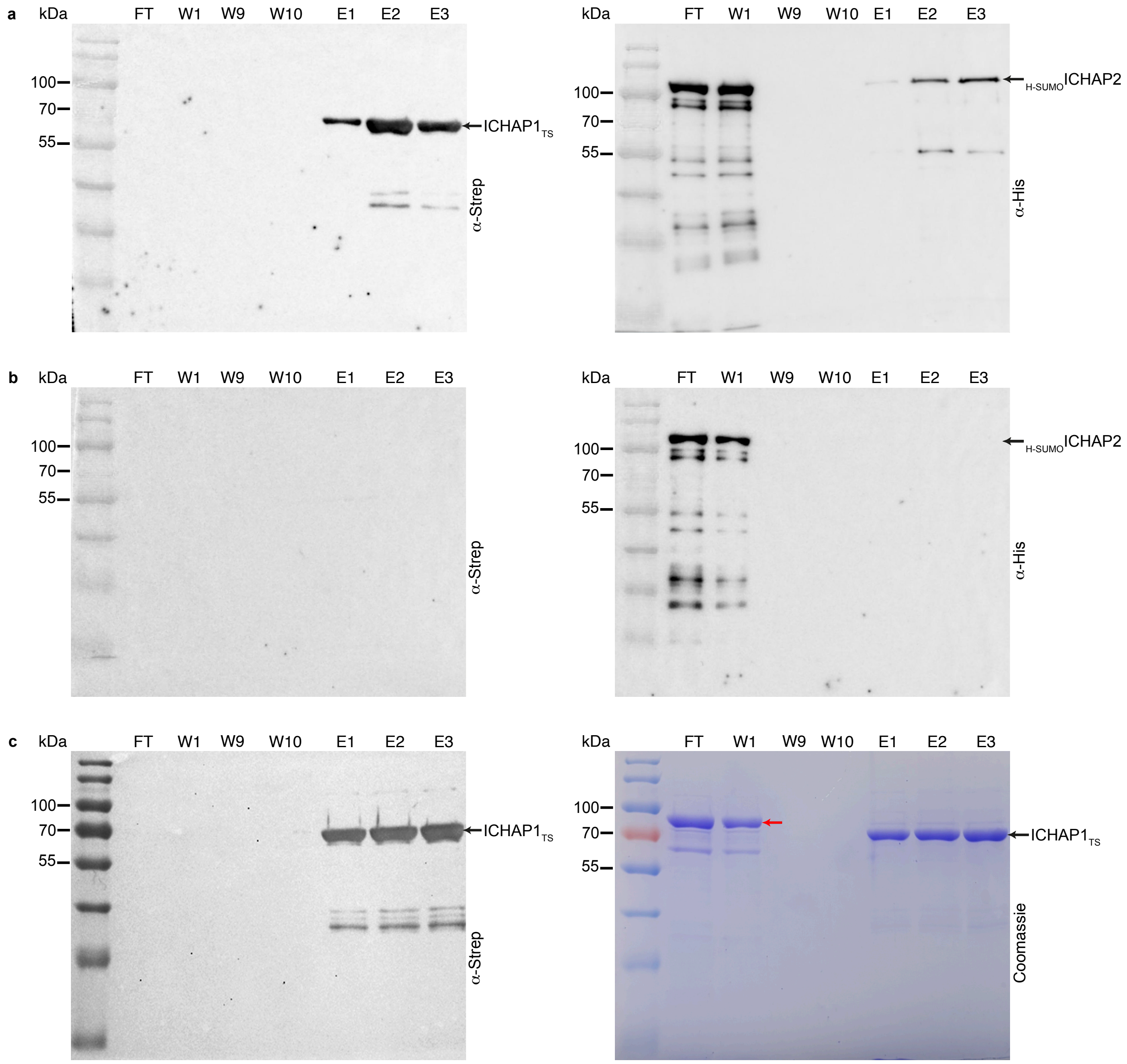
