## Supporting Table S1 for "*Medicago truncatula* Iron-chaperone 1 (ICHAP1) is required for symbiotic nitrogen fixation"

**Table S1. Putative KH-domain containing proteins expressed in *M. truncatula* roots and nodules.** Expression data (number of reads) have been obtained from the Symbimics database. ZI indicates nodule Zone I; ZIId, distal Zone II (region closer to ZI); ZIIp, proximal Zone II; IZ, interzone; and ZIII, Zone III.

| ***M. truncatula* gene** | **Root** | **Nodule** | **Nodule ZI** | **Nodule ZIId** | **Nodule ZIIp** | **Nodule IZ** | **Nodule ZIII** |
|---|---|---|---|---|---|---|---|
| *Medtr1g090773* | 2981.6 ± 205.6 | 2103.9 ± 213.2 | 314.6 ± 13.5 | 234.7 ± 11.9 | 146.6 ± 3.5 | 137.3 ± 8.7 | 167.9 ± 20.1 |
| *Medtr1g070380* | 1323.6 ± 34.3 | 1552.9 ± 74.3 | 109.5 ± 2.3 | 115.3 ± 12 | 96.8 ± 16.5 | 80.4 ± 2.2 | 115.3 ± 16.7 |
| *Medtr1g016970* | 1513.6 ± 40.9 | 2023.2 ± 22.5 | 166.9 ± 3.6 | 214.8 ± 9.6 | 146.3 ± 60.7 | 253.7 ± 17.3 | 280.4 ± 34.5 |
| *Medtr1g016760* | 2910.4 ± 323.5 | 3268.2 ± 175.2 | 139.6 ± 1.7 | 150.9 ± 13.1 | 167.6 ± 6.5 | 155.1 ± 16 | 245.7 ± 45.4 |
| *Medtr2g038020* | 856.4 ± 47.7 | 915.3 ± 6.4 | 186.1 ± 14.4 | 193.4 ± 7.7 | 224.6 ± 31 | 164.8 ± 15.3 | 124.7 ± 17.8 |
| *Medtr2g009650* | 764 ± 32.5 | 1043.6 ± 30 | 96.2 ± 2.9 | 113.9 ± 10 | 149.4 ± 24.3 | 181.7 ± 10.1 | 194.5 ± 29.4 |
| *Medtr2g019090* | 1054.2 ± 43.7 | 1605 ± 19.5 | 108.7 ± 8.4 | 101.1 ± 4.8 | 172 ± 30 | 175.2 ± 18.4 | 373.2 ± 17.6 |
| *Medtr3g104270* | 1892.3 ± 31.7 | 1886.9 ± 87 | 373.2 ± 11 | 320.6 ± 22.8 | 226.3 ± 13.7 | 114.5 ± 11.7 | 234.7 ± 28.3 |
| *Medtr3g091800* | 411.9 ± 55.3 | 538.9 ± 18.6 | 185.5 ± 22 | 182.7 ± 4.4 | 251.7 ± 29.4 | 251.6 ± 14.6 | 197.3 ± 17.3 |
| *Medtr3g110650* | 968.6 ± 89.6 | 1315.5 ± 29 | 126.7 ± 5.5 | 142.4 ± 19.3 | 102.8 ± 14.4 | 126.5 ± 9 | 83.8 ± 4.2 |
| *Medtr3g104280* | 1892.3 ± 31.7 | 1886.9 ± 87 | 373.2 ± 11 | 320.6 ± 22.8 | 226.3 ± 13.7 | 114.5 ± 11.7 | 234.7 ± 28.3 |
| *Medtr4g093650* | 213.4 ± 97.7 | 437.6 ± 11.6 | 32.2 ± 1.4 | 29.6 ± 2.3 | 23.7 ± 9.4 | 23.6 ± 7.6 | 19.1 ± 5.4 |
| *Medtr4g008080* | 596.3 ± 47.2 | 708.9 ± 14.4 | 71.1 ± 7.2 | 72.4 ± 4.9 | 52.1 ± 13.3 | 36.4 ± 2.1 | 53.7 ± 12.2 |
| *Medtr4g127380* | 895.7 ± 66 | 1346.1 ± 31.6 | 176.4 ± 5.4 | 190.6 ± 3.8 | 144.9 ± 12.6 | 148.4 ± 20 | 178.2 ± 7.2 |
| *Medtr5g008420* | 2645.5 ± 425 | 2822.3 ± 122.8 | 339.6 ± 5.5 | 319.1 ± 46.6 | 144.4 ± 46.4 | 88.8 ± 6.9 | 113.2 ± 18.2 |
| *Medtr5g031720* | 1633.6 ± 89.4 | 1763.1 ± 30.5 | 44.2 ± 4.7 | 53.2 ± 4.1 | 47.5 ± 20.8 | 93 ± 18.9 | 123.8 ± 6.8 |
| *Medtr5g049770* | 1633.6 ± 89.4 | 1763.1 ± 30.5 | 44.2 ± 4.7 | 53.2 ± 4.1 | 47.5 ± 20.8 | 93 ± 18.9 | 123.8 ± 6.8 |
| *Medtr6g012630* | 3354.2 ± 83.2 | 2938.2 ± 120.4 | 230.2 ± 10.1 | 222.3 ± 10.1 | 259.7 ± 100.6 | 239.1 ± 7.8 | 263 ± 27.8 |
| *Medtr7g115340* | 36.4 ± 4 | 32 ± 2.3 | 2.2 ± 0.9 | 5.8 ± 0.9 | 6.2 ± 3.6 | 4.7 ± 2.4 | 3.3 ± 1.7 |
| *Medtr7g011080* | 2365.7 ± 70.7 | 2856.9 ± 145.9 | 209.6 3.6 | 195.6 ± 7.1 | 172.5 ± 46.2 | 199.6 ± 23.1 | 316.8 ± 9.1 |
| *Medtr7g082290* | 52.9 ± 3.6 | 29 ± 10.6 | - | 0.5 ± 0.5 | - | - | - |
| *Medtr7g080630* | 390 ± 9.7 | 391.3 ± 9.2 | 71.7 ± 5.5 | 60.4 ± 5.3 | 53 ± 14 | 53.9 ± 6.6 | 57.7 ± 8.6 |
