## Supporting Table S2 for "*Medicago truncatula* Iron-chaperone 1 (ICHAP1) is required for symbiotic nitrogen fixation"

**Supplemental Table S2: Primers used in this study.**

| Primer | Sequence | Use |
|---|---|---|
| MY2Hb NRAMP1 N-tag FW | AAGAGGTGGTATGCACAGATCAGCTTTGTCGACG GTATCGATGGCACATCAAGAAGTGG | NRAMP1 expression in pBT3N MY2H vector |
| MY2Hb NRAMP1 N-tag RV | CGTGACATAACTAATTACATGACCTATTAA GATCTGACGTTCACAACCTAGGTCCCTCT | NRAMP1 expression in pBT3N MY2H vector |
| MY2Hb VTL8 N-tag FW | AAGAGGTGGTATGCACAGATCAGCTTTGTCG ACGGTATCGATGGCCGTTGGTACAATAT | VTL8 expression in pBT3N MY2H vector |
| MY2Hb VTL8 N-tag RV | CGTGACATAACTAATTACATGACCTATTAAGA TCTGACGTTCAAATTTCCAAATCCAAA | VTL8 expression in pBT3N MY2H vector |
| MY2Hb FPN2 N-tag FW | AAGAGGTGGTATGCACAGATCAGCTTTGTCG ACGGTATCGATGGATGAAAAAGAAGTTT | FPN2 expression in pBT3N MY2H vector |
| MY2Hb FPN2 N-tag RV | CGTGACATAACTAATTACATGACCTATTAAGA TCTGACGTTCATGCAGAAGAATTAATC | FPN2 expression in pBT3N MY2H vector |
| MY2Hb FPN2 C-tag FW | GCATTGCTGCTAAAGAAGAAGGGGTATCTTTG GATAAAAGAATGGATGAAAAAGAAGTTT | FPN2 expression in pAMBV MY2H vector |
| MY2Hb FPN2 C-tag RV | TGGAGGGATCCCCCCCGACATGGTCGACGGT ATCGATAAGCTGCAGAAGAATTAATCGAC | FPN2 expression in pAMBV MY2H vector |
| MY2Hp ICHAP1 C-tag FW | GATCCAAGCAGTGGTATCAACGCAGAGTGGC CATTATGGCTGGTCAAAGAACAT | ICHAP1 expression in pPR3C MY2H vector |
| MY2Hp ICHAP1 C-tag RV | CGTAATCTGGAACATCGTATGGGTACATAT CGATAAGCTTGTAACCATGGTTTCTCCTC | ICHAP1 expression in pPR3C MY2H vector |
| MY2Hp ICHAP2 C-tag FW | GATCCAAGCAGTGGTATCAACGCAGAGTGGC CATTATGGGTGAGAATGGTAAAA | ICHAP2 expression in pPR3C MY2H vector |
| MY2Hp ICHAP2 C-tag RV | CGTAATCTGGAACATCGTATGGGTACATATCG ATAAGCTTTGTGGCCATTATGAATGCC | ICHAP2 expression in pPR3C MY2H vector |
| NRAMP1 N-BiFC GW FW | GGGGACAAGTTTGTACAAAAAAGCAGGCTTCATG GCACATCAAGAAGTGGAT | Cloning of NRAMP1 in pNXGW BiFC vector |
| NRAMP1 N-BiFC GW RV | GGGGACCACTTTGTACAAGAAAGCTGGGTCTCAC AACCTAGGTCCCTCTGGTTG | Cloning of NRAMP1 in pNXGW BiFC vector |
| VTL8 N-BiFC GW FW | GGGGACAAGTTTGTACAAAAAAGCAGGCTTCATG GCCGTTGGTACAATATGT | Cloning of VTL8 in pNXGW BiFC vector |
| VTL8 N-BiFC GW RV | GGGGACCACTTTGTACAAGAAAGCTGGGTCTCAA ATTTCCAAATCCAAACCACT | Cloning of VTL8 in pNXGW BiFC vector |
| ICHAP1 C-BiFC GW FW | GGGGACAAGTTTGTACAAAAAAGCAGGCTTCATG GCTGGTCAAAGAACATCC | Cloning of ICHAP1 in pXCGW BiFC vector |
| ICHAP1 C-BiFC GW RV | GGGGACCACTTTGTACAAGAAAGCTGGGTCGTAA CCATGGTTTCTCCTCCG | Cloning of ICHAP1 in pXCGW BiFC vector |
| ICHAP2 C-BiFC GW FW | GGGGACAAGTTTGTACAAAAAAGCAGGCTTCATG GGTGAGAATGGTAAAAGA | Cloning of ICHAP2 in pXCGW BiFC vector |
| ICHAP2 C-BiFC GW RV | GGGGACCACTTTGTACAAGAAAGCTGGGTCTGTG GCCATTATGAATGCCTG | Cloning of ICHAP2 in pXCGW BiFC vector |
| 5 MtUb v4qF | ATTCTTCACATGCGGCGATTTAC | qRT-PCR of ubiquitin carboxylterminal hydrolase |
| 3 MtUb v4qR | TTTCTCATTTGCTTTTGGTGTTGG | qRT-PCR of ubiquitin carboxylterminal hydrolase |
| 5 ICHAP1 ATG + 665 qRT FW | GGCTCGGCTTCATGACAACCCT | qRT-PCR of ICHAP1 |

| | | |
|---|---|---|
| 3 ICHAP1 ATG + 750 qRT RV | GAGCCCCACCATGAGAACCCAC | qRT-PCR of ICHAP1 |
| 5in-fuspET16b ICHAP1 | AGGAGATATACCATGGCTGGTCAAAGAACATCC | Cloning of ICHAP1 in modified pET16b vector |
| 3in-fus ICHAP1::TEV::TS | ATGACTCCAGCCCATACCTTGAAAATAAAGATTTTCGTAACCATGGTTTCTCCTCC | Cloning of ICHAP1 in modified pET16b vector |
| 5in-fus C-TS | ATGGGCTGGAGTCATCCTCAGTTTGAGAAA | Cloning of ICHAP1 in modified pET16b vector |
| 3in-fus TS-stop::pET16b | GTTAGCAGCCGGATCCTCATTTTTCAAATTGTGGATGTGA | Cloning of ICHAP1 in modified pET16b vector |
| 5in-fus His-SUMO-ICHAP2 | CTTCCAATCCAATATTATGGTAAAAGAAATCGTCAGC | Cloning of ICHAP2 in pET His6x-SUMO vector |
| 3in-fus His-SUMO-ICHAP2 | CACCAGGTGGCGCGCCTCATTTTTCAAATTGTGGATG | Cloning of ICHAP2 in pET His6x-SUMO vector |
| ICHAP1p -2000 GW FW | GGGGACAAGTTTGTACAAAAAAGCAGGCTTCCATAATGTTTATGCACCCGTA | Cloning of ICHAP1p in pGWB3/ ICHAP1p + genomic ICHAP1 in pGWB13 and in pGWB1 |
| ICHAP1p -21 GW RV | GGGGACCACTTTGTACAAGAAAGCTGGGTCTGTTTTGTGTGAGAACTGTGT | Cloning of ICHAP1p in pGWB3 |
| ICHAP1ATG +3980 (w/o STOP) GW RV | GGGGACCACTTTGTACAAGAAAGCTGGGTCGTAACCATGGTTTCTCCTCCG | Cloning of ICHAP1p + genomic ICHAP1 in pGWB13 and in pGWB1 |
| ICHAP1 (Phusion promoter – CDS) FW | ACACAGTTCTCACACAAAACAATGGCTGGTCAAAGAACATCCT | Cloning of ICHAP1p + genomic ICHAP1 in pGWB1 |
| ICHAP1 (Phusion promoter – CDS) RV | AGGATGTTCTTTGACCAGCCATTGTTTTGTGTGAGAACTGTGT | ICHAP1p + genomic ICHAP1 in pGWB1 |
